# Mechanophenotyping of 3D Multicellular Clusters using Displacement Arrays of Rendered Tractions

**DOI:** 10.1101/809871

**Authors:** Susan E. Leggett, Mohak Patel, Thomas M. Valentin, Lena Gamboa, Amanda S. Khoo, Evelyn Kendall Williams, Christian Franck, Ian Y. Wong

## Abstract

Epithelial tissues mechanically deform the surrounding extracellular matrix during embryonic development, wound repair, and tumor invasion. *Ex vivo* measurements of such multicellular tractions within three-dimensional (3D) biomaterials could elucidate collective dissemination during disease progression, and enable preclinical testing of targeted anti-migration therapies. However, past 3D traction measurements have been low throughput due to the challenges of imaging and analyzing information-rich 3D material deformations. Here, we demonstrate a method to profile multicellular clusters in a 96-well plate format based on spatially heterogeneous contractile, protrusive, and circumferential tractions. As a case study, we profile multicellular clusters across varying states of the epithelial-mesenchymal transition, revealing a successive loss of protrusive and circumferential tractions, as well as the formation of localized contractile tractions with elongated cluster morphologies. These cluster phenotypes were biochemically perturbed using drugs, biasing towards traction signatures of different epithelial or mesenchymal states. This higher-throughput analysis is promising to systematically interrogate and perturb aberrant mechanobiology, which could be utilized with human patient samples to guide personalized therapies.

## Introduction

Collective mechanical interactions between epithelial cells and 3D extracellular matrix (ECM) shape embryonic tissue development, and their dysregulation can drive cancer progression or other disease states^1^. In particular, the epithelial-mesenchymal transition (EMT) is associated with weakened cell-cell junctions and increased cell-matrix adhesions, driving tissue disorganization and dissemination^2^. Remarkably, multicellular clusters cultured *ex vivo* in compliant 3D hydrogels can exhibit tissue-like form and function^3^, representing a promising preclinical testbed for higher-throughput drug discovery and development^4^. However, existing assays for 3D mechanophenotyping have been limited to a few cells per experiment due to the need for high-resolution optics, labor-intensive image processing, as well as complex readouts of force generation^5^. Rapid biophysical characterization of multicellular clusters in a 3D matrix could enable direct characterization and perturbation of disease state in human patient samples, informing personalized and predictive therapies^6^.

Traction force microscopy (TFM) resolves cell-generated deformations based on the motion of fluorescent tracer particles embedded within a compliant material^7^. Early TFM studies measured cell-cell and cell-matrix interactions of multicellular epithelial clusters on planar 2D substrates^8–13^. More recently, TFM has been extended to 3D hydrogels, but has primarily focused on individual cells^14–24^. Notably, epithelial cell lines exhibit more isotropic, spatially uniform tractions, while mesenchymal cell lines exhibit highly anisotropic tractions localized at the leading and trailing edge^14–16, 18, 20, 22, 23^. Nevertheles, it remains a formidable challenge to visualize 3D cell morphologies and tractions, particularly in a scalable experimental format^25^. Most traditional TFM approaches require relatively high-resolution data on both cellular morphologies and material deformations, which poses significant experimental, workflow and computational challenges. This issue is further exacerbated by also requiring knowledge of the underlying material properties to compute cellular tractions, which continues to be a significant challenge for fibrous and biologically remodelled extracellular materials^7^.

Here, we demonstrate a high content method to profile the spatially heterogeneous matrix deformations of multicellular clusters in a standard 96 well plate format. We show that clusters exhibit collective tractions with distinct spatial signatures, which we visualize by tracking hundreds of thousands of embedded tracer particles to recover information-rich material displacement fields. This Displacement Arrays of Rendered Tractions (DART) analysis was validated by inducing EMT in mammary epithelial cells cultured within a composite 3D hydrogel of silk fibroin and collagen I. Remarkably, we find that the progression from epithelial to mesenchymal states is associated with a successive loss of protrusive and circumferential tractions, as well as the formation of highly localized contractile tractions. Indeed, these emergent behaviors cannot be resolved using conventional spatially averaged TFM metrics developed for individual cells. We further perturb these cluster mechanophenotypes towards more mesenchymal or epithelial states using drugs that stabilize microtubules (e.g. Taxol) or inhibit epidermal growth factor receptor signaling (e.g. Gefitinib). Since this approach does not rely on high-resolution 3D object detection, lower numerical aperture air objectives can be used to rapidly image a multiwell plate for higher-throughput volumetric image acquisition and analysis. This implementation of 3D TFM on a standardized platform could enable preclinical screening of human organoids to inform drug development and personalized therapies.

## Displacement Arrays of Rendered Tractions (DART)

We designed a new kinematics based approach called displacement array of rendered tractions (DART) metrics for characterizing the cell-induced matrix deformations without relying on constitutive material model for the 3D biomaterial. We previously demonstrated mean deformation metrics (MDM) to compute average values associated with the deformation fields of individual cells^21^. However, we observed in these experiments that multicellular clusters applied spatially heterogeneous deformation fields over large volumes. Thus, the corresponding MDM regressed to null values due to spatial averaging. Moreover, this spatial averaging resulted in a loss of biologically interesting information associated with highly localized signatures in the cell-induced deformation field, making MDM unsuitable for analyzing collective deformations.

To compute the DART metrics, the particle displacement (*u*_*i*_) at positions (*x*_*i*_) were interpolated on to regularized grid points (*x*_*grid*_) in each image with a general spacing of 12 voxels in the *x*, *y* and *z* directions, which provided good trade-off in terms of spatial resolution and computational throughput. Let *u*_*grid*_ be the displacement vector at the regularized grid points *x*_*grid*_. A linear scattered interpolation scheme was utilized. The DART metrics compute kinematic quantities from the local deformation field of the cell cluster. Thus, only the displacement *u*_*grid*_ at points *x*_*grid*_ within at distance *d* = 25 *µ*m from the cell cluster surface were considered in computing the DART metrics, which was sufficient to capture most of the displacement data around each cluster. The displacements *u*_*grid*_ at the points outside the search region were set to be zero. At each of the grid points *x*_*grid*_, a unit vector *n*_*grid*_ originating from the centroid of the cell cluster (*r*_*o*_) to the *x*_*grid*_ was determined. Using the normal vector, the displacement *u*_*grid*_ was then decomposed into radial 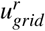 and hoop 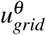 displacement components as,

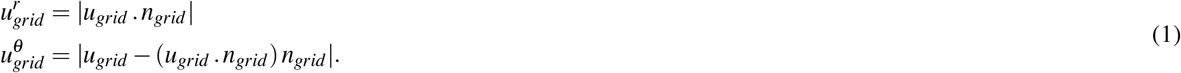

From 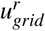 and 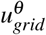, the regions where cells applied significant protrusive, contractile and circumferential deformations were determined. These regions are found by binarizing the 3D matrix of 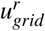 and 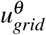 using thresholding operation with *u*_significant_ as

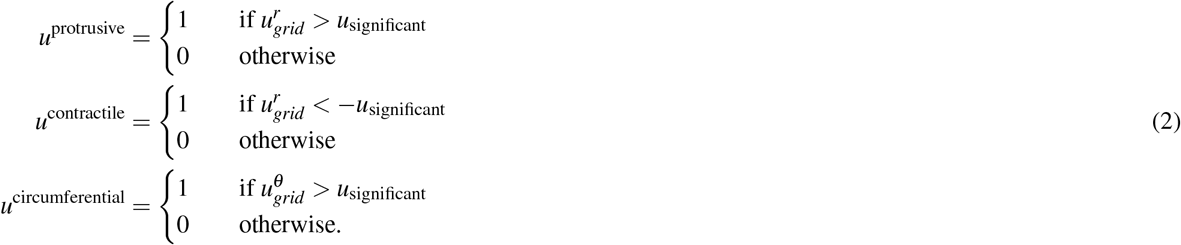

The level of *u*_significant_ can have significant implications on the computed DART metrics, and was chosen to be at least five times above the intrinsic displacement noise floor and to produce an optimally even distribution of displacement slices in the DART control groups. To remove spurious white voxels in *u*^protrusive^, *u*^contractile^ and *u*^circumferential^ produced from noise in particle tracking, these 3D matrix were filtered to remove connected white components smaller than our 12 voxels grid point spacing corresponding to a volume of 106.5 *µ*m^3^.

From *u*^protrusive^, *u*^contractile^ and *u*^circumferential^ the relative localized regions where cells apply protrusive, contractile and circumferential deformation is computed through DART. Here, the computation of contractile DART is described, but protrusive and circumferential DARTs are computed analogously. The volume about the center of the cell cluster in *u*^contractile^ is divided into 16 equal sub-volumes. The volume around each cell in *u*^contractile^ is first divided into two, bottom and top, hemispheres. Each hemisphere is further divided into 8 equal, *π*/4 apart, regions. Thus, each “slice” in the DART corresponds to a volumetric region in the real space around the cell cluster. The outer and inner slices in the DART board correspond to the region in the lower (*θ*_*el*_ ≥ *π*) and upper (*θ*_*el*_ < *π*) hemispheres of the *u*^contractile^ region around each cell cluster. A slice is considered to be contractile if the corresponding region in *u*^contractile^ has at least one white voxel signifying that the cell cluster applies significant contractile deformation in that region. The contractile quantification within each DART slice is binary. From the contractile DART, the number of contractile displacement slices is used as a metric to quantify how localized the contractile deformation is that the cell cluster applies. Similarly, the number of protrusive and circumferential displacement slices for each cell cluster was computed.

As the final part of our DART approach, a phenotype classifier was built to classify multi-cellular clusters into epithelial, mesenchymal and transitory phenotypes using the DecisionTreeClassifier class from the Python Scikit library, a commonly used machine learning tool. For classification, the decision tree utilized the number of contractile, protrusive and circumferential displacement slices and shape anisotropy factor of a multi-cellular cluster. The decision tree classifier had a maximum tree depth of 5 and a minimum number of leaf samples of 5. The decision tree model was trained with low tree depth and high minimum number of leaf samples to prevent training data overfitting. The decision tree was trained on all multi-cellular clusters used in the analysis of the control DMSO treatment condition. The accuracy of the decision tree on the training data is evaluated using the normalized confusion matrix. A normalized confusion matrix **C** is defined such that **C**_*i j*_ is the proportion of the observation known to be in the group *i* and classified in the group *j*. The confusion matrix of a perfect classifier is equal to the identity matrix.

## Results

### DART Visualizes Spatially Heterogeneous Displacement Fields

Collective cell-matrix interactions were characterized by embedding mammary epithelial cells (MCF-10A) in a composite hydrogel consisting of 7.5 mg/mL silk fibroin and 1 mg/mL collagen I^26^ on a 96 well plate (Fig. 1a). These hydrogels had an elastic modulus of 600 Pa and characteristic pore size of 2 *µ*m (**Fig. S1**), which maintained epithelial cells as spherical clusters (“acini”) but was also permissive for local dissemination (**Fig. S2**). These MCF-10A Snail-ER cells were stably transfected to overexpress green fluorescent protein (GFP) in the cytoplasm, as well as an inducible estrogen receptor construct for controlled EMT through the Snail transcription factor^27, 28^. Cluster-induced deformations of the hydrogel were visualized using our topological particle tracking algorithm to map the displacement of 80,000-100,000 tracer particles around each cluster^29^, after which the clusters were lysed to define a reference state. This case study focused on clusters cultured for 7 days, which exhibited distinct morphologies (**Fig. S2**).

**Figure 1.**
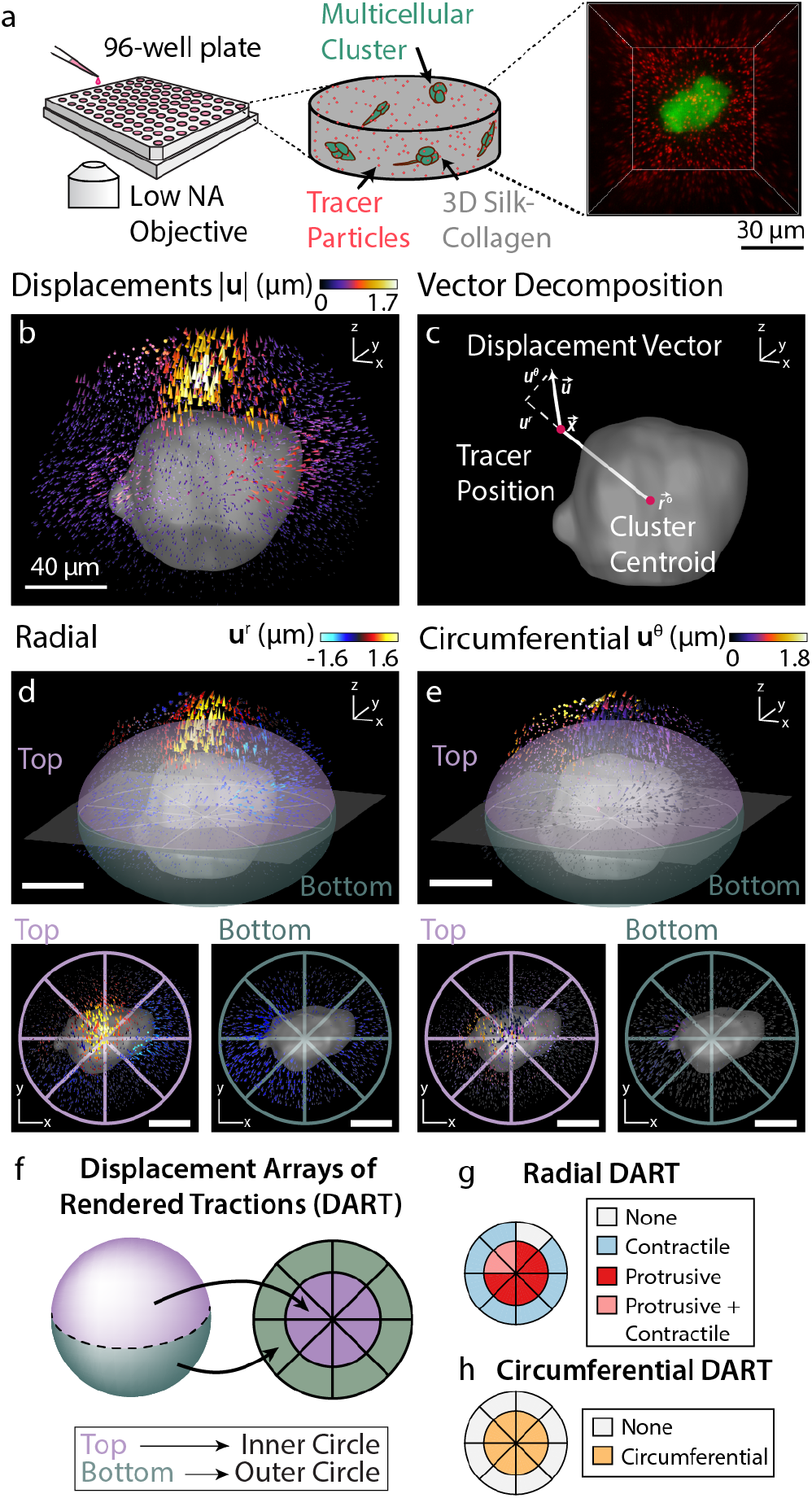
Schematic illustration of the experimental setup and displacement array of rendered tractions (DART) metrics. **(a)** Experimental setup for high-throughput imaging to measure cell-induced matrix deformations. Multicellular clusters were grown inside a silk-collagen matrix with embedded 1 *µ*m red fluorescent tracer particles in a 96 well plate setup. To achieve high-throughput imaging, clusters were imaged using a spinning disk confocal microscopy with a low numerical aperture air objective. **(b)** 3D cell-induced matrix deformations recovered by directly tracking tracer particles. **(c)** Matrix displacements 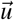 were decomposed into radial (*u*^*r*^) and circumferential (*u*^*θ*^) components about the center of the cluster. **(d)** *top*: Radial displacements *u*^*r*^ of the matrix around the cell cluster. The surrounding volume is partitioned into top and bottom hemispheres, which were projected onto a two-dimensional representation in the *lower left* and *lower right* respectively. **(e)** *top*: Circumferential displacements *u*^*θ*^ of the matrix around the cell cluster. The surrounding volume were partitioned into top and bottom hemispheres, which are again projected onto a two-dimensional representation in the *lower left* and *lower right* **(f)** Schematic mapping of three-dimensional displacement fields onto a two-dimensional DART representation. **(g)** Protrusive or contractile displacements with magnitude larger than *u*_significant_ in the surrounding volume were represented by protrusive or contractile slices in the radial DART metrics. The top and bottom hemispheres map to the inner and outer slice of the DART respectively. **(h)** Circumferential DART computed from **e** analogous to the method described in **g**.

Multicellular clusters typically deformed the surrounding matrix in a spatially heterogeneous manner. For instance, a representative cluster exhibited inward (“contractile”) tractions around the periphery, but outward (“protrusive”) tractions near the top (Fig. 1b). Although visually pronounced, this heterogeneity is often averaged out in conventional TFM or kinematic metrics based on mean deformations around individual cells^14, 21^ **(Fig. S3)**. In order to profile these spatial patterns, the displacement vector of a given tracer particle 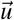 due to matrix deformation was decomposed into radial *u*^*r*^ and circumferential components *u*^*θ*^ relative to the center of the cluster (Fig. 1c). These discrete particle displacements were then interpolated onto a regularized grid in order to optimize computational throughput while maintaining adequate spatial resolution. Next, the volume around each cluster was subdivided into sixteen equal sub-volumes, with eight regions of equal volume in top and bottom hemispheres, respectively (Fig. 1d,e). For ease of visualization, we developed the Displacement Arrays of Rendered Tractions (DART) analyses to map these 3D deformations onto a simpler, 2D representation. In each DART, the inner region of eight equal slices corresponds to the eight subvolumes in the top hemisphere, while the outer ring corresponds to the bottom hemisphere (Fig. 1f,g). If the radial particle displacement within a given region exceeded a critical threshold (*u*_*significant*_) of 0.4 *µ*m (**Fig. S4**), it was denoted as contractile, protrusive, or both (Fig. 1f). Similarly, if the circumferential displacement within a given region exceeded a certain threshold of 0.4 *µ*m, it was also noted (Fig. 1g). This DART displacement threshold, *u*_significant_ was chosen to be at least five times above the intrinsic displacement noise floor and to produce an optimally even distribution of displacement slices in the DART control groups (see **Fig. S4**). Our displacement noise floor obtained with our topology-based particle tracking algorithm was 20.5 nm^29^. Finally, the spatial heterogeneity of tractions was then quantified from the number of regions which exceeded the radial or circumferential displacement threshold.

### Epithelial, Transitory, and Mesenchymal Clusters Exhibit Distinct Tractions and Morphologies

As a case study, three experimental conditions were characterized with multicellular clusters representing epithelial, transitory, and mesenchymal mechanophenotypes **(Fig. S5)**. MCF-10A Snail-ER cells were embedded as single cells within silk-collagen hydrogels and then imaged after 7 days. First, MCF-10A cells were maintained in an epithelial mechanophenotype by culturing with 0.05% dimethyl sulfoxide (DMSO), matching the concentration used to resuspend drug compounds. Epithelial cells organized into compact, roughly spherical multicellular acini (Fig. 2a). These clusters exerted localized protrusive and some circumferential deformations relative to the reference state after lysing, while the spatial distribution of contractile deformations varied across clusters (Fig. 2b,c). Second, MCF-10A cells were induced to a transitory (EMT) mechanophenotype by Snail induction with 500 nM 4-hydroxytamoxifen (OHT) after embedding in the hydrogel. These transitory clusters exhibited significant protrusions (Fig. 2d), analogous to the budding outgrowths associated with branching morphogenesis. Moreover, these clusters exhibited uniformly distributed contractile deformations across the periphery with some circumferential deformations, but minimal protrusive deformations (Fig. 2e,f). Third, MCF-10A cells were pre-induced into a mesenchymal mechanophenotype by sustained treatment with 500 nM 4-hydroxytamoxifen (OHT) for 72 hours before embedding into the hydrogel^27^. These clusters were highly elongated and spindle-like, with slightly decreased sizes due to slower proliferation after EMT (Fig. 2g). Mesenchymal clusters exhibited highly localized contractile deformations at only a few locations, consistent with front/back polarity, as well as minimal protrusive or circumferential deformations (Fig. 2h,i).

**Figure 2.**
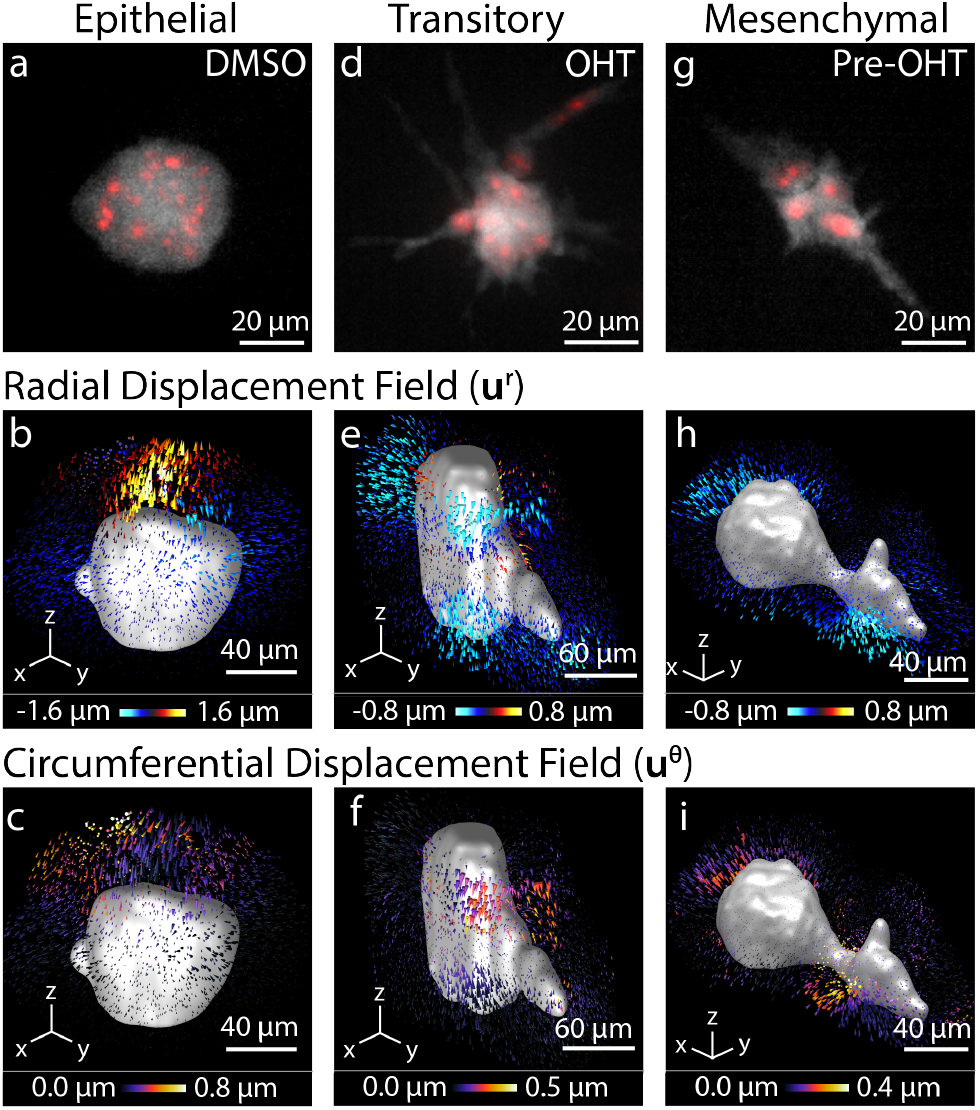
Representative mechanophenotypes corresponding to epithelial, transitory, and mesenchymal states. **(a,d,g)** Morphology of the cell clusters. The cytoplasm of the cell cluster is shown in gray (GFP) and the nucleus of the cells are shown in red (mCherry-H2B). **(b,e,h)** 3D radial displacements, *u*^*r*^, about the center of a cell cluster shown for a representative epithelial, mesenchymal, or transitory cell cluster. **(c,f,i)** Corresponding 3D circumferential displacements, *u*^*θ*^, about the center of a cell cluster shown for a representative epithelial, mesenchymal, or transitory cell cluster.

Representative DART analyses for clusters with epithelial, transitory, and mesenchymal mechanophenotypes captured these qualitative trends, although there existed appreciable heterogeneity across clusters. Epithelial and transitory clusters exerted spatially distributed contractile displacements over many subvolumes, while mesenchymal clusters exerted localized contractile displacements over fewer subvolumes (Fig. 3a,b). Next, epithelial clusters applied a significant number of protrusive displacements, which were considerably lower for transitory and mesenchymal clusters (Fig. 3a,c). Moreover, epithelial and transitory clusters also exerted significant circumferential displacements, which were much lower for mesenchymal clusters (Fig. 3d, **S6a,b,c)**. Finally, epithelial and transitory clusters exhibited relatively compact morphologies based on shape anisotropy, while mesenchymal clusters were very elongated (Fig. 3e) These distinct cluster mechanophenotypes were compared based on their number of protrusive and contractile slices, revealing that epithelial clusters typically exhibited spatially distributed protrusive deformations, while the distribution of contractile deformations varied (Fig. 3f). In comparison, transitory clusters typically exhibited localized protrusive deformations but highly distributed contractile deformations around the periphery (Fig. 3f). Lastly, mesenchymal clusters exhibit minimal protrusive deformations with a varying distribution of contractile deformations (Fig. 3f).

**Figure 3.**
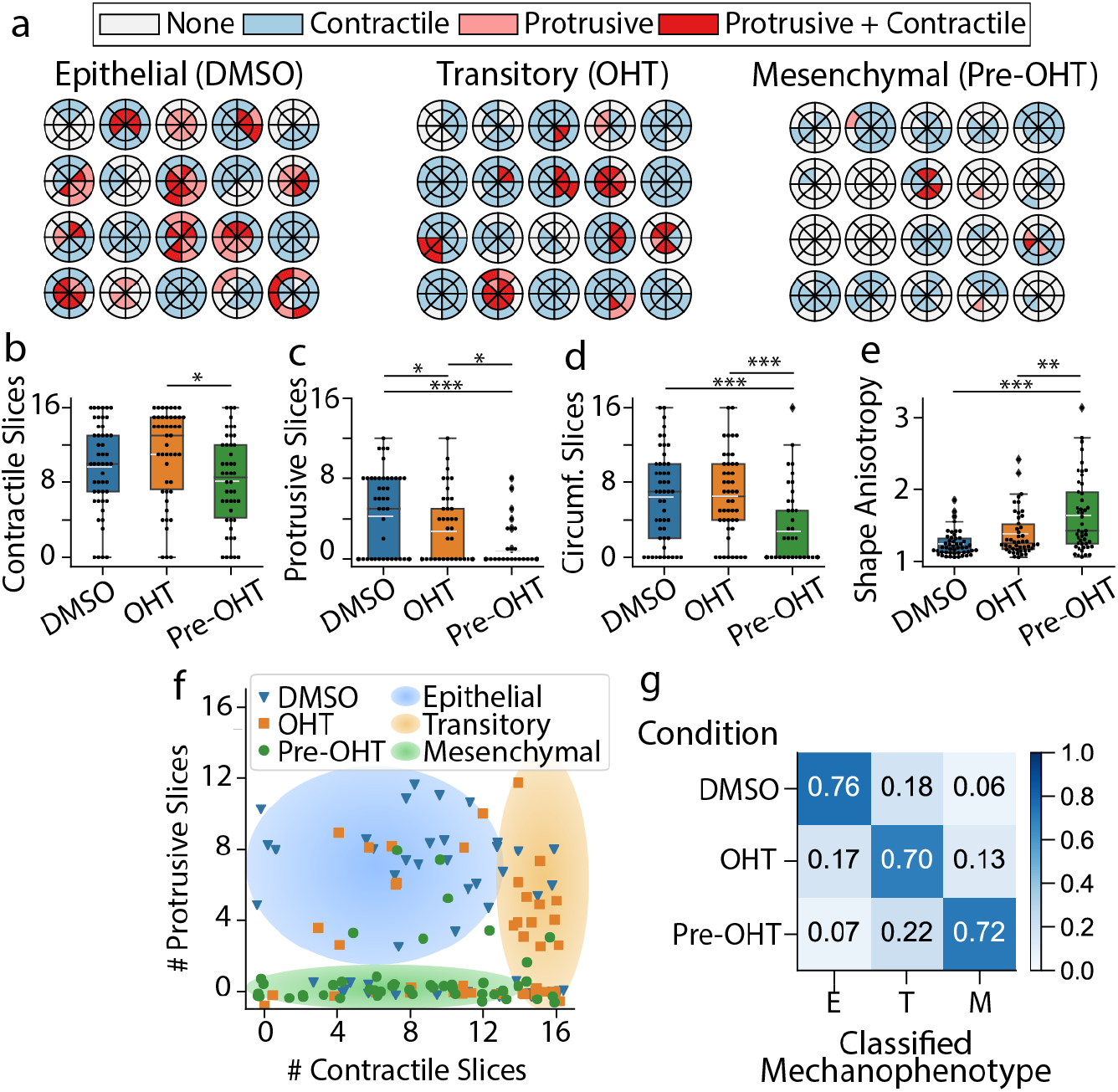
**(a)** Radial DARTS for twenty randomly selected clusters treated with DMSO, OHT, or pre-treated with OHT. **(b)** Boxplot of the number of contractile displacement slices. **(c)** Boxplot of the number of protrusive displacement slices. **(d)** Boxplot of the number of circumferential displacement slices. **(e)** Boxplot for cell morphology shape anisotropy. The white dashed line in the boxplot represents the mean value. The symbols *, **, and *** represent *p* values of *<*0.05, *<*0.01 and *<*0.001 respectively. **(f)** Scatter plot of raw data points for number of protrusive and contractile displacement slices (*P*_*s*_ and *C*_*s*_) for DMSO treated, OHT treated, or OHT pre-treated clusters. Jitter has been applied to the true positions of the raw data points to avoid points occlusion. The shaded regions are a guide to the eye to emphasize groupings of epithelial, transitory, or mesenchymal mechanophenotypes. **(g)** Normalized confusion matrix for the decision tree classifier utilizing *C*_*s*_, *P*_*s*_, number of circumferential displacement slices (*θ*_*s*_) and shape anisotropy (*SA*) metrics on the training data.

Using a decision tree, a commonly employed predictive classifier in computer science and machine learning, these three distinct mechanophenotypes were profiled based on contractile, protrusive, and circumferential deformations, as well as shape anisotropy (**Supporting Information**). Briefly, this analysis classified clusters based on a threshold number of protrusive slices (4.5), which was then refined based on shape anisotropy, number of circumferential slices, and contractile slices **(Fig. S7)**. We assessed and quantified the specificity and predictive capability of our decision tree via the standard machine learning approach of using a confusion matrix. The classification via the confusion matrix showed 70% agreement between experimental condition and predicted mechanophenotype (epithelial, transitory, mesenchymal), as shown in the on-diagonal entries of the confusion matrix (Fig. 3g). Moreover, 15-20% of clusters in each experimental condition were classified in an adjacent state (i.e. epithelial clusters classified as transitory, transitory clusters classified as epithelial or mesenchymal, etc.), corresponding to the neighboring off-diagonal entries of the confusion matrix (Fig. 3g). This classification may be be attributed to biological heterogeneity, since MCF-10A cells can spontaneously undergo EMT, and EMT induction kinetics exhibit some variability^30^. Nevertheless, classification across very dissimilar mechanophenotypes was relatively infrequent at 6-7% (i.e. epithelial clusters classified as mesenchymal, or vice-versa), corresponding to the entries at the top right and bottom left corners of the confusion matrix (Fig. 3g). Thus, epithelial, transitory, and mesenchymal clusters exhibit distinct morphologies and patterns of contractile, protrusive, and circumferential traction, which represent a characteristic “traction signature” or mechanophenotype.

### Microtubule Stabilization with Taxol Enhances Protrusions and Localized Contractility

Cluster mechanophenotypes were perturbed by sublethal treatment with the microtubule stabilizing agent Taxol, which can induce EMT^30^. After 7 days of treatment with a sublethal dose of Taxol (4 nM) and 0.05% DMSO, clusters exhibited partially elongated morphologies reminiscent of the transitory mechanophenotype with OHT treatment only **(Fig. S8abc)**. Similarly, treatment with 4 nM Taxol and 500 nM OHT resulted in highly elongated morphologies reminiscent of the mesenchymal mechanophenotype after pretreatment of 500 nM OHT only **(Fig. S8def)**. Interestingly, pretreatment with 500 nM OHT and 4 nM Taxol resulted in unique cluster morphologies with slender, neuronal-like extensions **(Fig. S8ghi)**. DART analysis of the cluster-induced matrix deformations corroborated these qualitative observations (Fig. 4a, **S6def)**. For instance, Taxol and DMSO-treated clusters exhibited more contractile, fewer protrusive, and more circumferential slices, analogous to the transitory cluster mechanophenotype (Fig. 4b,c,d), **S6d**. In comparison, Taxol and OHT-treated clusters exhibited fewer contractile and circumferential slices, analogous to the mesenchymal cluster mechanophenotype (Fig. 4b,d), **S6e**. Finally, Taxol and OHT-pretreated clusters exhibited fewer contractile and circumferential slices and elevated shape anisotropy, also consistent with the mesenchymal cluster mechanophenotype (Fig. 4b,d,e), **S6f**. Interestingly, Taxol and OHT treated or pretreated clusters exhibited more protrusive slices than the comparable transitory and mesenchymal mechanophenotypes. These behaviors may be attributed to the stabilizing action of Taxol on outward cytoskeletal protrusions, which cannot retract as easily since microtubule depolymerization is inhibited^31^. These trends were apparent on a plot of the contractile and protrusive slices per cluster, since Taxol and DMSO-treated clusters were shifted rightward with more contractile slices (Fig. 4f) relative to the epithelial mechanophenotype in DMSO only (Fig. 3f). Moreover, Taxol and OHT treated or pretreated clusters were shifted upwards with more protrusive slices (Fig. 4f) relative to transitory and mesenchymal mechanophenotype with OHT treatment or pretreatment only (Fig. 3f). As a consequence, 35% of Taxol and DMSO treated clusters were classified as a transitory mechanophenotype (Fig. 3g). Similarly, 37% of Taxol and OHT treated transitory clusters were classifed as a mesenchymal mechanophenotype (Fig. 3g). Finally, 60% of Taxol and OHT pretreated mesenchymal clusters were classified as a mesenchymal (OHT only) mechanophenotype (Fig. 3g). It should be noted that a significant percentage (24-43%) of Taxol-treated clusters were classified as epithelial mechanophenotype, likely due to the increased number of protrusive slices (Fig. 3g). Thus, Taxol treatment biases clusters towards more transitory and mesenchymal mechanophenotypes by redistributing contractile and circumferential tractions, although protrusive tractions are also aberrantly enhanced relative to the previous experiments without Taxol.

**Figure 4.**
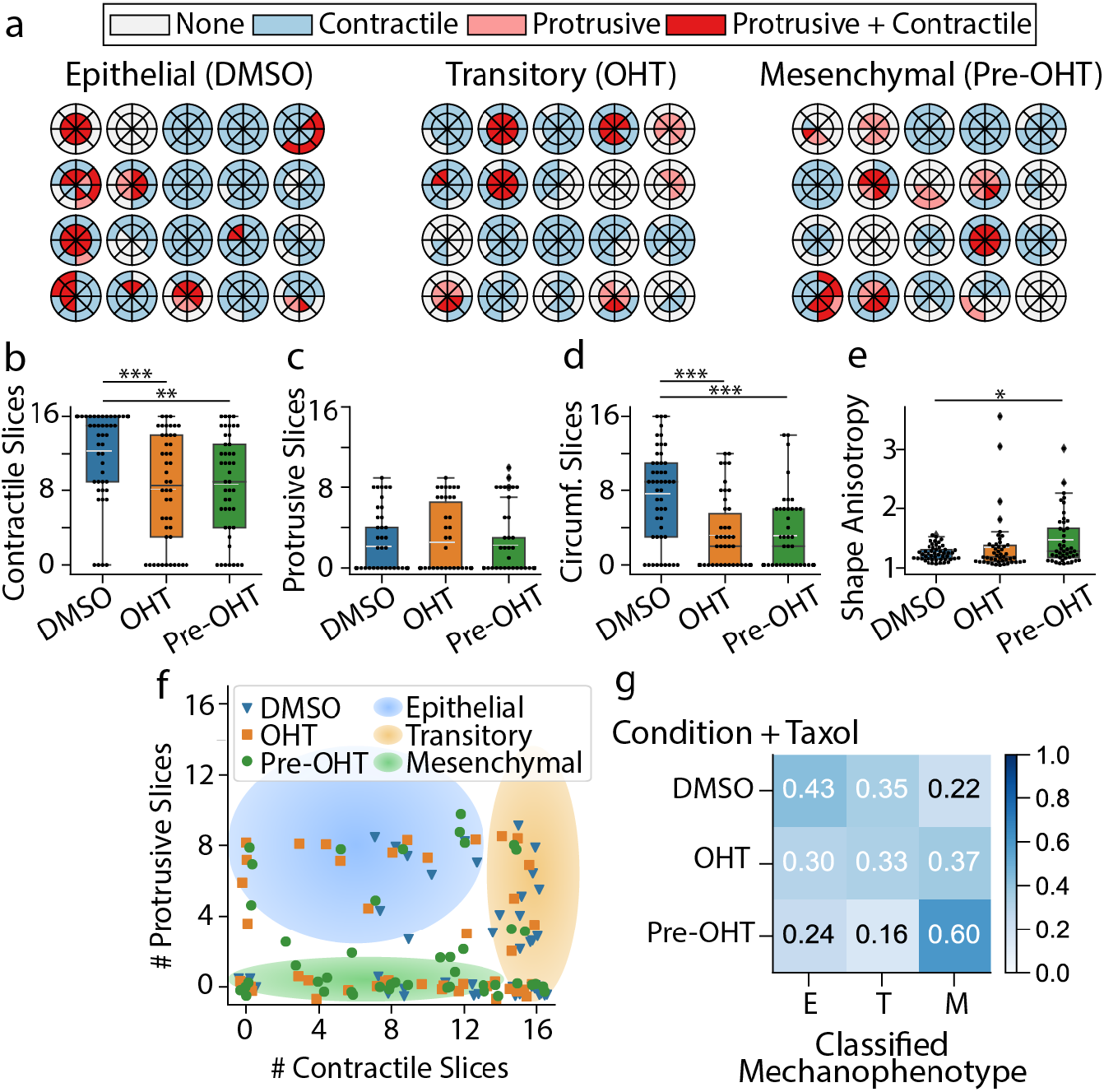
**(a)** Radial DARTS for twenty randomly selected clusters treated with 4 nM Taxol combined with DMSO, OHT, or pre-treatment with OHT. **(b)** Boxplot of the number of contractile displacement slices. **(c)** Boxplot of the number of protrusive displacement slices. **(d)** Boxplot of the number of circumferential displacement slices. **(e)** Boxplot for cell morphology shape anisotropy. The white dashed line in the boxplot represents the mean value. The symbols *, **, and *** represent *p* values of *<*0.05, *<*0.01 and *<*0.001 respectively. **(f)** Scatter plot of raw data points for number of protrusive and contractile displacement slices (*P*_*s*_ and *C*_*s*_) for DMSO treated, OHT treated, or OHT pre-treated clusters. Jitter has been applied to the true positions of the raw data points to avoid points occlusion. The shaded regions are a guide to the eye to emphasize groupings of epithelial, transitory, or mesenchymal mechanophenotypes. **(g)** Normalized confusion matrix for the decision tree classifier utilizing *C*_*s*_, *P*_*s*_, number of circumferential displacement slices (*θ*_*s*_) and shape anisotropy (*SA*) metrics on the training data.

### Epidermal Growth Factor Receptor Inhibition with Gefitinib Increases Heterogeneity of Transitory Clusters

Lastly, cluster mechanophenotypes were perturbed by treatment with the epidermal growth factor receptor (EGFR) inhibitor Gefitinib, which ordinarily inhibits proliferation in EGF-dependent MCF-10A cells. Nevertheless, EMT induction is associated with decreased sensitivity to such EGFR (tyrosine kinase) inhibitors^32^. After 7 days treatment with 500 nM Gefitinib (also a sublethal dose), DMSO-treated clusters exhibited compact morphologies consistent with the epithelial mechanophenotype **(Fig. S9abc)**. In comparison, Gefitinib and OHT-treated clusters exhibited both compact and elongated morphologies, reminiscent of epithelial and mesenchymal phenotypes **(Fig. S9def)**. Finally, Gefitinib and OHT-pretreated clusters exhibited highly elongated morphologies, reminiscent of mesenchymal phenotypes **(Fig. S9ghi)**. DART analysis revealed that Gefitinib and DMSO treated clusters exhibited more contractile, protrusive, and circumferential slices with low shape anisotropy (Fig. 5a-e, **S6g)**, consistent with a epithelial mechanophenotype. However, Gefitinib and OHT-treated clusters exhibited large variations in protrusive and circumferential slices relative to the transitory (OHT only) cluster mechanophenotype (Fig. 5a,c,d), **S6h**. Finally, Gefitinib and OHT-pretreated clusters exhibited fewer contractile, protrusive, and circumferential slices with slightly increased shape anisotropy, analogous to a mesenchymal mechanophenotype (Fig. 5a-e), **S6i**. A plot of contractile and protrusive slices per cluster showed that Gefitinib and DMSO-treated clusters were located in the top left region associated with the epithelial (DMSO only) mechanophenotype. Similarly, Gefitinib and OHT-pretreated clusters were located at the bottom of the plot, associated with the mesenchymal (OHT-pretreatment only) mechanophenotype. In comparison, the Gefitinib and OHT treated clusters were widely dispersed towards the top and bottom of the plot (Fig. 5f). The decision tree analysis revealed that 73% and 64% of Gefitinib with DMSO or OHT-pretreated clusters were classified as epithelial (DMSO only) and mesenchymal (OHT pretreated) mechanophenotypes, respectively (Fig. 5f). Nevertheless, Gefitnib and OHT-treated clusters were mostly classified as either epithelial (42%) or mesenchymal (42%) mechanophenotypes, with relatively few clusters classified as transitory clusters (16%). Thus, Gefitinib treatment did not significantly affect the DMSO or OHT-pretreated cluster, which retained epithelial or mesenchymal mechanophenotype, respectively. However, Gefitnib and OHT treatment resulted in a mixed population of mostly epithelial and mesenchymal mechanophenotypes, with considerably fewer transitory clusters.

**Figure 5.**
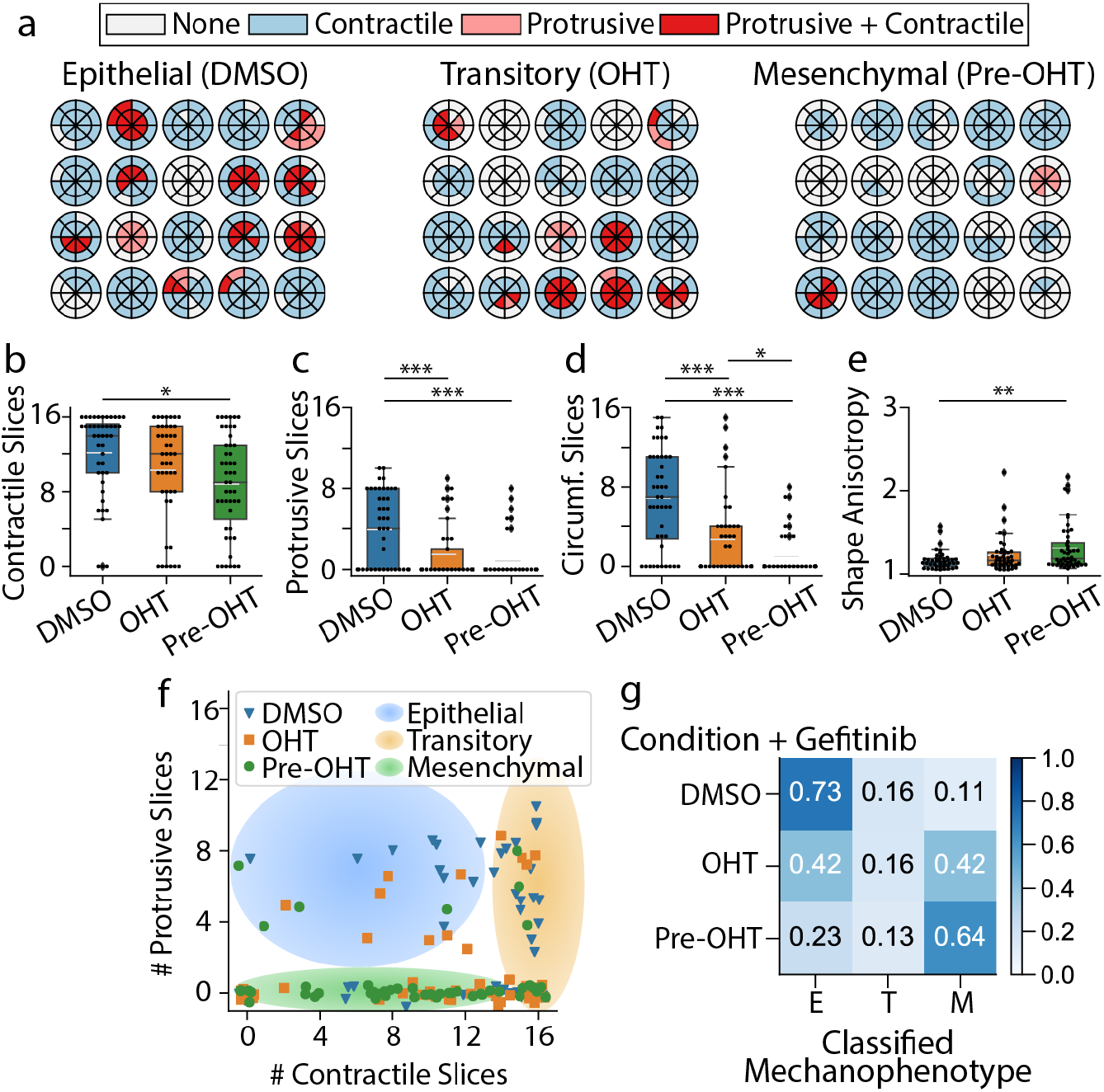
**(a)** Radial DARTS for twenty randomly selected clusters treated with 500 nM Gefitinib combined with DMSO, OHT, or pre-treatment with OHT. **(b)** Boxplot of the number of contractile displacement slices. **(c)** Boxplot of the number of protrusive displacement slices. **(d)** Boxplot of the number of circumferential displacement slices. **(e)** Boxplot for cell morphology shape anisotropy. The white dashed line in the boxplot represents the mean value. The symbols *, **, and *** represent *p* values of *<*0.05, *<*0.01 and *<*0.001 respectively. **(f)** Scatter plot of raw data points for number of protrusive and contractile displacement slices (*P*_*s*_ and *C*_*s*_) for DMSO treated, OHT treated, or OHT pre-treated clusters. Jitter has been applied to the true positions of the raw data points to avoid points occlusion. The shaded regions are a guide to the eye to emphasize groupings of epithelial, transitory, or mesenchymal mechanophenotypes. **(g)** Normalized confusion matrix for the decision tree classifier utilizing *C*_*s*_, *P*_*s*_, number of circumferential displacement slices (*θ*_*s*_) and shape anisotropy (*SA*) metrics on the training data.

## Discussion and Conclusion

The DART analysis established here profiles multicellular cluster mechanophenotypes based on local displacements of tracer particles embedded in a 3D matrix, as well as cluster morphology. By combining these measurements into a machine learning decision tree structure, perturbations and transitions from one phenotype to another (e.g., epithelial to mesenchymal) as a function of drug treatment can be quantified in an automated fashion. This approach leverages previous developments in the machine learning communities of reducing large, multivariable spatiotemporal data sets and binning them into a more intuitive scalar metric output.

These purely kinematic measurements do not rely on mechanical or microstructural properties of the matrix, which to-date remain poorly understood for biologically-derived materials such as our silk fibroin-collagen I hydrogel. Such fibrous materials exhibit highly nonlinear and nonaffine responses, which may not be easily captured using classical continuum description based on bulk rheological measurements^33^. As a result, DART in its current form reports cellular traction signatures using displacement field measures. As appropriate constitutive descriptions of the extracellular matrix become available, DART could be augmented with mechanical tractions, stresses and forces. Similarly, DART’s predictive power could be further refined by imaging other subcellular features or biomarkers, which could be incorporated into the decision tree. Finally, it should also be noted that the reference state was defined at the completion of the experiment by lysing clusters and observing the subsequent relaxation of the matrix. This reference state does not account for irreversible, inelastic deformations of the matrix, which may occur due to ECM remodeling or deposition. In future work, complementary measurements using optical tweezers^20, 23^ or direct imaging of fibers^18^ could also be incorporated to address how multicellular clusters are affected by changes in the local stiffness of the surrounding matrix.

Our classification of multicellular cluster mechanophenotype directly accounts for local spatial heterogeneity in the local 3D matrix deformation patterns. Indeed, we find distinct patterns to these spatial deformations that are sufficient to classify 70% of cluster mechanophenotypes within a given experimental condition based on DMSO treatment, OHT treatment, or OHT pretreatment. It should be noted that some clusters were classified with a different mechanophenotype than would be expected from their experimental conditions. For instance, 17% of clusters treated with OHT were classified as an epithelial mechanophenotype, rather than transitory. Moreover, 18% of clusters treated with DMSO were classified as a transitory mechanophenotype rather than epithelial. These differences likely reflect inter-cluster heterogeneity, since EMT may occur slower or faster in distinct clusters, resulting in distinct mechanophenotypes at a given snapshot in time. Interestingly, circumferential displacements were significant for epithelial and transitory clusters but less pronounced for mesenchymal clusters. This traction field may occur due to the previously observed orbiting motions observed for epithelial clusters^34, 35^, which are less likely to occur for highly asymmetric cluster morphologies with weakened cell-cell junctions. These differences are also observed after treatment with EMT inducing or suppressing drugs, indicative of differences in drug sensitivity. Although computationally expensive, we envision that time-lapse imaging of clusters and associated tractions could further reveal how clusters interconvert between mechanophenotypes.

In conclusion, DART establishes a quantitative and scalable framework to profile the heterogeneous matrix deformations of 3D multicellular clusters. Our analyses reveal that collective tractions are spatially non-uniform and cannot be captured by existing analyses for individual cells that assume isotropic behavior. We show that epithelial mechanophenotypes typically apply protrusive, circumferential and contractile matrix deformations, while transitory mechanophenotypes after EMT exhibit mostly contractile and circumferential deformations that are widely distributed, and mesenchymal clusters exhibit localized contractility in only a few locations. We perturb these behaviors using the microtubule stabilizer Taxol, which biases towards transitory or mesenchymal mechanophenotype while also enhancing protrusions. In comparison, the EGFR inhibitor Gefitinib drives clusters towards either epithelial or mesenchymal mechanophenotype in OHT treated conditions, but has minimal effect on epithelial (DMSO) or mesenchymal (OHT pretreated) mechanophenotype. Thus, DART captures both intra- and inter-heterogeneity of multicellular clusters in response to biochemical stimulation. We envision that DART can be implemented with a wide variety of 3D biomaterials at a 96 well plate scale or beyond, enabling higher throughput mechanophenotyping of organoids in 3D culture, including preclinical testing of human patient samples with personalized treatments.

## Methods

### Cell Culture and Matrix Preparation

MCF-10A mammary epithelial cells stably transfected with an inducible Snail expression construct fused to an estrogen receptor response element were a gift from D.A. Haber (Massachusetts General Hospital)^27^. This cell line also overexpressed fluorescent proteins in the nucleus (mCherry-H2B) and cytoplasm (GFP) for live cell tracking. MCF-10A cells were routinely cultured in growth media consisting of DMEM/F12 HEPES buffer (Fisher 11330057) supplemented with 5% horse serum (Fisher 16050122), 20 ng/ml Animal-Free Recombinant Human Epidermal Growth Factor (EGF; PeproTech AF-100-15), 0.5 mg/mL hydrocortisone (Sigma H0888), 100 ng/mL cholera toxin (Sigma C8052), 10 *µ*g/mL Insulin from bovine pancreas (Sigma I1882), and 1% Penicillin-Streptomycin (Fisher MT-30-002-CI).

Silk fibroin solution was extracted and purified from silkworm (*B. mori*) cocoons (Treenway Silks, Lakewood, CO), as previously described^26^. Composite silk-collagen hydrogels were prepared through sonication-induced gelation initiation of silk fibroin, followed by the addition and neutralization of collagen I from rat tail tendon (Corning 354249) to achieve a final hydrogel containing 7.5 mg/mL silk fibroin and 1 mg/mL collagen I (see **SI Appendix** for details). Briefly, silk fibroin was mixed into media and then sonicated, 1N sodium hydroxide was added in to achieve a final pH of 7.4, collagen 1 was mixed in well, followed by the addition of 5% 1*µ*m fluorescent carboxylate-modified beads (Fluospheres, red 580/605) and lastly, a single-cell suspension in media was mixed in to yield 120,000 cells/mL.

Cells were embedded in 3D silk-collagen hydrogels with three experimental conditions: 1) Cells were cultured in 0.05% dimethyl sulfoxide (DMSO, the solvent used to suspend OHT), for 72h in 2D culture then embedded in 3D hydrogels with DMSO treatment. This condition serves as the negative control and defined the “epithelial” mechanophenotype. 2) Cells were treated with 0.05% DMSO for 72h in 2D culture, as in (1), then embedded in 3D hydrogels with 500 nM of 4-hydroxytamoxifen (“OHT”, Sigma H7904). This condition induces Snail expression through an estrogen receptor construct^27^ and defined the “transitory” mechanophenotype. 3) Cells were treated with OHT for 72h in 2D culture to induce EMT then embedded in 3D hydrogels with sustained OHT treatment to maintain Snail expression as a positive control, which defined the “mesenchymal” mechanophenotype.

### Confocal Microscopy and Image Analysis

Multicellular clusters and corresponding matrix displacements were imaged after 7 days of culure using a Nikon Eclipse TiE fluorescence microscope with spinning-disk confocal head (Crest Optics X-light V2), with light-guide coupled solid state illumination system (Lumencor Spectra-X3), sCMOS camera (Andor Neo), 20x Plan Apo objective (NA 0.75), GFP/FITC Filter Set (Chroma 49002), TRITC/DSRed Filter Set (Chroma 49004). For the duration of time-lapse imaging, cells were maintained in a humidified environmental chamber at 37 °C, 5% CO_2_. For matrix displacement measurements, NIS Elements was used for automated image acquisition with z-steps of 0.6 *µ*m from the bottom of the well to a height of 75 *µ*m under consistent exposure times, camera gain/gamma control, and aperture. Images were acquired at 4 h intervals over a large number of wells (*n* ≈ 48 for each experiment) for a total of 16 h. In these experiments, cell cytoplasm was imaged in GFP and beads in the RFP channel. At the end of time-lapse imaging, a reference state for the gel was obtained by lysing the cells within the hydrogels via sodium dodecyl sulfate.

### Measuring 3D Cell-induced Deformations via Topology-based Particle Tracking

We utilized our previously developed topology-based particle tracking (TPT) algorithm to reconstruct the cell-induced 3D displacement fields by tracking individual fluorescent polystyrene microspheres (1 *µ*m) embedded as fiduciary markers in the silk-collagen matrix. The combination of LED illumination-based spinning disk confocal microscopy and low numerical aperture objective for low-cost, high-throughput imaging of the 96-well plate setup resulted in diminished signal to noise volumetric images. A custom image segmentation and filtering routine was developed to allow precise and accurate localization and tracking using TPT for low numerical aperture confocal imaging stacks (see SI for details).

### Cell Cluster Surface Segmentation

The 3D cell cluster surface was segmented from volumetric images of fluorescently labeled cytoplasm intensities (GFP channel). As a first step, the raw volumetric images were filtered using a median filter with 3 × 3 × 3 voxel window to remove shot noise. Following the images were filtered with a 3D Gaussian filter with *σ* = 2.5. The images were then binarized using adaptive image thresholding based on the local mean intensity (first-order statistics) in the neighborhood of each voxel. The sensitivity for the adaptive thresholding was manually set for each image to segment the cell clusters from the background appropriately. From the binary images, the small connected components having a total number of voxels less than 8000 voxels were set to an intensity value of 0 in the binary images. Morphological operations were performed to remove holes in the binary images^36^. The volumes of the segmented binary images were increased by 1.6 *µ*m through a distance transform. Due to the large noise near the top and bottom of the volume, all the voxels in the top and bottom 8 z-slices were set to 0 in the binarized images. The 3D triangulated cell cluster surface was computed from the binary images using MATLAB’s isosurface estimation at a target voxel value of 0.5. The triangulated cell cluster surface mesh was smoothed using accurate curvature flow smoothing^37^.

### Statistical Analysis

Experiments were repeated 3 times (external replicates), and a total of at least 40 clusters were analyzed per experimental condition. To compare DART and shape metrics across phenotype conditions, one–way repeated measures analysis of variance (One Way RM ANOVA) was used to check if the treatment data satisfied the Shapiro-Wilk normality test with *p <* 0.05. If the treatment data failed the Shapiro-Wilk normality test, Friedman repeated measures analysis of variance on ranks (RM ANOVA on ranks) for comparing treatment differences was used. For all pairwise multiple comparisons Holm-Sidak post-hoc test was used. The differences were considered to be statistically significant if *p <* 0.05. The statistical tests were performed using SigmaPlot 12.0.

In the figures, we use boxplots to visualize the distribution for each metrics. As per convention, boxplots show the data median, first quartile, third quartile, and data outliers marker through minimum and maximum values. The dots on the boxplot plot show the raw data values. The white dashed lines on the boxplots show the mean value. In the graphs, a statistically significant difference between two treatments is shown by a line connecting their boxplot with an annotation for the *p*-value. The symbols ∗, ∗∗, and ∗∗∗ denote *p* values of *<* 0.05, *<* 0.01 and *<* 0.001 respectively.

## Acknowledgements

We thank C.M. Nelson and J.S. Reichner for careful readings, as well as D.A. Haber for the inducible MCF-10A cell lines. This work was supported by NIH Grants T32ES007272, P30GM110759, R21CA212932, and Brown University (Karen T. Romer Undergraduate Research and Teaching Award, and Start-Up Funds).

## Author contributions statement

S.E.L., M.P., C.F., and I.Y.W. designed research; S.E.L. performed experiments; M.P. contributed new analytic tools; M.P. and S.E.L. analyzed data with feedback from C.F., and I.Y.W.; S.E.L., T.M.V., L.G., A.S.K., and E.K.W. prepared and characterized materials; S.E.L., M.P., C.F., and I.Y.W. wrote the paper with feedback from all authors

