## Supplementary Information for "Mechanophenotyping of 3D Multicellular Clusters using Displacement Arrays of Rendered Tractions"

### Methods

**Cell Culture.** MCF-10A mammary epithelial cells stably transfected with an inducible Snail expression construct fused to an estrogen receptor response element were a gift from D.A. Haber (Massachusetts General Hospital) (1). This cell line also overexpressed fluorescent proteins in the nucleus (mCherry-H2B) and cytoplasm (GFP) for live cell tracking. MCF-10A cells were routinely cultured in growth media consisting of DMEM/F12 HEPES buffer (Fisher 11330057) supplemented with 5% horse serum (Fisher 16050122), 20 ng/ml Animal-Free Recombinant Human Epidermal Growth Factor (EGF; PeproTech AF-100-15), 0.5 mg/mL hydrocortisone (Sigma H0888), 100 ng/mL cholera toxin (Sigma C8052), 10 $\mu$g/mL Insulin from bovine pancreas (Sigma I1882), and 1% Penicillin-Streptomycin (Fisher MT-30-002-CI) (2). Snail expression was induced by supplementing the media with 500 nM of 4-hydroxytamoxifen ("OHT", Sigma I1882) (1) in dimethyl sulfoxide.

**Silk Fibroin Extraction.** Silk fibroin solution was extracted and purified from silkworm (*B. mori*) cocoons (Treenway Silks, Lakewood, CO), as previously described (3). Briefly, cocoons were cut into ≈1 cm$^2$ pieces. To remove sericin, 5 g of silk cocoons were added to 2 L of boiling water containing 4.24 g sodium carbonate. "Purified water" ASTM Type II Deionized Distilled water (442-88, MilliporeSigma) was used for all steps. Silk cocoons were boiled for 20 minutes and gently but continuously mixed using a metal spatula. After boiling, the silk cocoons were immediately placed into 2 L of purified water at room temperature for 20 min. The water was then regularly replaced with purified water, for a total of 3 water changes. The cocoons were then set to dry overnight, after which silk was teased apart manually using tweezers, with care to remove clumps. A 20% (w/v) solution was then made with silk fibers and 60°C LiBr, with a final volume of 15 mL. The silk and LiBr solution was left to sit at 60°C until all of the silk fibers had been solubilized. The solubilized silk solution was then dialyzed against 1 L of purified water for at least 48 hrs., changing the water a total of 6 times. Silk was then centrifuged at 4°C, 5,000 RPM for 20 minutes, and decanted to remove any large debris from cocoons. This process was repeated for two centrifugation steps. The purified silk solution was then stored at 4°C to limit $\beta$-sheet formation and crosslinking. The concentration of the purified silk solution was determined by pipetting 1 mL of the purified silk solution into a weigh boat and placing it in the 60°C oven to dry overnight. The dried silk film was then weighed to calculate the w/v concentration.

**Silk-Collagen Composite Hydrogel Preparation.** In order to prepare composite silk-collagen hydrogels for cell culture experiments, a series of initial optimization steps must first be carried out to ensure 1) physiological conditions will be met upon embedding cells in the 3D microenvironment by adjusting the pH to 7.4 and 2) a homogeneous matrix will be obtained from the interpenetrating network of silk and collagen, which is verified by the dispersion of fluorescently labeled, carboxylate-modified microspheres ("beads", 1 $\mu$m red fluorescent 580/605 carboxylate-modified FluoSpheres, ThermoScientific). First, a pH array was conducted for every new batch of silk fibroin, collagen I, or prepared media since these all affect the pH of the resulting hydrogel. Growth media without phenol red was used for preparation of all hydrogels to reduce background during fluorescence imaging. Briefly, based on the desired final concentration of components (7.5 mg/mL silk fibroin, 1 mg/mL collagen I, +/-5% beads and 1 mL total volume) the volume of media, SF, COL, and beads were determined and then added to a 2 mL microtube (Axygen MCT2000) in order, while on ice. The mixture was homogenized with five-pulse vortices at mid-intensity after the addition of each component. Next, the pH of the resulting mixture was measured, recorded, and adjusted to 7.4 using 1N sodium hydroxide (NaOH). Generally, 1-2 $\mu$L of 1N NaOH were required to adjust the pH of the mixture, with some batch to batch variability.

With the amount of NaOH required for a final hydrogel pH of 7.4 determined, the volume of media required was appropriately recalculated for subsequent steps. Next, a sonication array was performed to determine the amount of time required to induce silk fibroin $\beta$-sheet formation such that upon the addition of collagen I and subsequent incubation of the composite hydrogel, a homogeneous matrix will be formed with well-dispersed collagen I and tethered labeled fluorescencent beads. Note that this procedure was modified from our previous publication, which optimized the sonication time based on the gelation of silk-only hydrogels (3). To do this, beads were first sonicated in a water bath (power level 4) for at least 1 hour to break up any aggregates. A tissue culture treated 96 well CellCarrier imaging plate (6005550, PerkinElmer) was then placed in the incubator (37°C, 5% $CO_2$) to be pre-warmed while hydrogel components were prepared. Next, all steps were performed using aseptic technique in a biosafety cabinet and reagents were kept cold (4°C) at all times using ice and/or an iceblock for microtubes: 1) The appropriate amount of media was added to a 2mL microtube, followed by the addition of SF to obtain a final concentration of 7.5 mg/mL SF for the 1 mL final volume. 2) The mixture was homogenized with five-pulse vortices at mid-intensity, which was repeated for the addition of every subsequent component. 3) The silk fibroin/media mixture was sonicated using the sonicator Model 120 Sonic Dismembrator, with programmed set time (tested from 2-10 s) and 30% power. 4) Collagen I (1 mg/mL final), beads (5% final), and 1N NaOH were added in order, with mixing in between as described. 5) 60 $\mu$L of the final 1 mL solution was then pipetted into individual wells of the pre-warmed 96-well plate for several technical replicates (at least $n = 4$). 5) The plate was placed back into the incubator to allow for the gelation of the hydrogel to occur. 6) Steps 1-5 were repeated for all sonication times to be tested in the array. After 1 h of incubation, the hydrogels were gently topped off with 120 $\mu$L of phenol red free growth media to prevent their drying out. Hydrogels were incubated overnight and imaged the next day to visualize the dispersion of beads within the matrix. For newly prepared batches of silk fibroin, a 5 s sonication time could be consistently and reproducibly used.

**Unconfined Compression Stress-Relaxation Compression Testing.** The elastic modulus and stress relaxation half-life were measured using unconfined compression testing (4) on an Instron 5944 MicroTester precision instrument equipped with Instron Bluehill software, a 5 N load cell, and Screw Side Action Grips rated to 500 N (cat. no. 2710-004). 3D printed custom compression platens were fabricated from PLA using a Makerbot Replicator 3D Printer. To ensure a smooth compression surface, a Fisherbrand Unbreakable 22 × 22 mm cover slips were adhered to each platen surface using Scotch removable poster tape (cat. no. 109). The cross-sectional area was calculated from side-view images before and after compression using a Logitech HD Pro Webcam C910. Compression data and images were analyzed in Matlab 2018.

Samples were prepared in 2 mL microcentrifuge tubes. After polymerization, the bottoms of the tubes were removed using a No. 10 scalpel. The samples were then gently pushed out of the tube and cut to a height of 0.25 in. (6.35 mm) in height. To ensure accurate imaging of the hydrogel cross-section, excess water was gently wicked away from the sample before and after the compression test using a kimwipe.

The unconfined compression stress relaxation protocol involved compressing the sample to 25% final compressive strain in increments of 5% strain at 8 mm/s. Between compression steps, the platens were held in place, and the sample was allowed to relax for 10 min. Due to the sensitivity of the load cell and the lack of a vibration isolation table for the Instron, the load response data was smoothed in Matlab using a span of 10.

The elastic modulus was calculated by determining the slope of the stress–strain curve for each sample. This was determined by fitting a linear regression to the compressive strain (mm/mm) and the residual stress at the end of each relaxation period. The stress relaxation half-life is defined as the point at which the stress had relaxed to 50% of the peak load. This was calculated from the final compression step for each sample (from 20 to 25% compressive strain). The load response data was max-min normalized where the maximum stress was determined as the peak stress at the start of the relaxation period, and the minimum stress was determined as the minimum stress at the end of the relaxation period. Curve-fitting was then performed using a two-termed exponential model that showed good agreement with the relaxation data. Using this model, the stress relaxation half-life time point was calculated from the half-maximum stress.

**Second Harmonic Generation Imaging of Silk-Collagen Composite Hydrogels.** Second harmonic generation (SHG) imaging was performed on hydrogels prepared within open polydimethylsiloxane (PDMS) wells bonded to a 6-well plate. Briefly, PDMS was well mixed in a 10:1 ratio of the base component to the curing agent, degassed in a vacuum chamber for 15 min, and then cured in an oven at 60 °C for 1 h. Wells were then punched out of the PDMS using an 8 mm in diameter biopsy punch. Silk-collagen precursor solution was prepared as described above, then 200 $\mu$L of this solution was pipetted into each well, hydrogels were incubated for 1h, topped off with media to prevent dehydration, and imaging was conducted after 24h of total incubation.

SHG imaging was then conducted using an Olympus FV1000-MPE multiphoton laser scanning microscope at the Brown University Leduc Bioimaging Facility. The microscope was equipped with a Mai Tai HP tunable laser (690–1020 nm), four non-descanned detectors (2 PMTs and 2 GaAsP detectors), an encoded Prior Z deck with a scanning stage, a 405/40 SHG filter cube allowing selection of a stimulation wavelength between 780 and 840 nm, and an Olympus 25× 1.05 NA XLPlan Multiphoton water objective. Images were acquired 150–450 $\mu$m from the top surface of the hydrogel (n=3 images per sample). The laser was tuned to 800 nm and 3 W, the laser scanning speed was set to 10 $\mu$s/pixel with a 4× line scan average and a size of 1600 × 1600 pixels (508.8 $\mu$m × 508.8 $\mu$m). The photomultiplier (HV) was set between 525 and 675 V with the Gain set to 2× and Offset set to 9%.

Collagen fiber length and density were analyzed as described previously (3). SHG images ($n$ = 3 per condition) were first cropped to an 800 x 800 pixel image to eliminate artifacts from non-uniform illumination. Images were then analyzed using CT-FIRE for Individual Fiber Extraction v2.0 software for Linux (5) to extract fiber angle, length, straightness, and width. CT-FIRE parameters were optimized for the best segmentation of visible fibers and visually checked by overlaying detected fibers over a contrast optimized image. Parameters were kept consistent across images except for the following ranges, optimized per image manually: thresh_im2: 90-100, xlinkbox: 2-12. If CT-FIRE segmentation was satisfactory, CT-FIRE .mat files were read into CurveAlign, which further analyzes fiber length, width, and density.

The segmented fibers obtained from CT-FIRE and CurveAlign are typically output as subdivided fiber segments and identified by a unique ID. The end-points of each fiber segment are connected together (by line segments) to visualize the the complete fiber. To quantify fiber density, the central ROI (800 px by 800 px) in each SHG image was further subdivided into square boxes and the number of distinct fibers within each box was computed by checking for the presence of at least one fiber segment endpoint within the box. The number of fibers per box was then calculated by counting the number of fibers with unique fiber ids per each box. For ease of visualization, the number of fibers per 50 x 50 pixel boxsize was converted into microns and divided by 2.53 scaling factor to yield an equivalent graph depicting the number of fibers per every $\mu$m. The mean number of fibers over all boxes was recorded at various box sizes (side lengths: 5, 10, 20, 40, 50, 80, 100, 160, 200, 400 and 800 pixels; 1 px = 0.318 $\mu$m). A relationship between box size (box area in $\mu m^2$) and number of fibers was established by constructing a cubic interpolant. To estimate fiber density (and mesh area) in the image, the area corresponding to a single fiber was approximated using the cubic interpolation function. Pore diameter was then calculated by dividing by $\pi$, taking the square root and multiplying by 2.

**EMT in 3D Silk-Collagen Hydrogels.** Cells were embedded in 3D silk-collagen hydrogels with pH adjusted to 7.4. Three main Snail induction conditions were examined to yield a range of epithelial-mesenchymal phenotypes: 1) Cells were treated with OHT for 72 h in 2D culture to induce EMT then embedded in 3D hydrogels with continued OHT treatment to maintain Snail expression. This condition is referred to as "Mesenchymal." 2) Cells were treated with a paired concentration of dimethyl sulfoxide (0.05% DMSO), the solvent used to suspend OHT, for 72 h in 2D culture then embedded in 3D hydrogels with DMSO treatment. This condition serves as the negative control and is referred to as "Epithelial." 3) Cells were treated with 0.05% DMSO for 7 2h in 2D culture, as in (2), then embedded in 3D hydrogels with OHT treatment. This condition serves to induce Snail expression in a 3D microenvironment compared to the rapid EMT induction in 2D culture (1), and is referred to as "Transitory."

To embed cells in 3D hydrogels, MCF-10A cells from separate flasks (+/- OHT) were dissociated from the tissue culture flask as single cell suspensions using Accumax (Fisher Scientific), spun down, resuspended in phenol red free growth medium, and counted with a Cellometer Auto 1000 Counter (Nexcelom Bioscience). Cells were then resuspended to a final concentration of 2 million cells/mL, such that the addition of 60 $\mu$L of cell suspension to each hydrogel microtube would yield a total of 120K cells/mL. Hydrogels were prepared as described in the previous section. Briefly, media (calculated volume less 60 $\mu$L to leave room for the addition of cells) was added to a 2 mL microtube on a 4°C tube rack. SF was added, the mixture was vortexed 5 times, sonicated for the optimal predetermined amount of time, COL was added, then NaOH, then +/- 5% beads. Note, for biological experiments beads were included to measure cell-induced matrix displacement fields, while in separate experiments, beads were excluded to allow for clear visualization of cell cytoplasm (GFP) and nuclear (RFP) fluorescence-based signal positions. Cells were then added to the hydrogel mixture as the final step, which was gently vortexed 5 times, and then 60 $\mu$L of the hydrogel-cell mixture was added to a pre-warmed 96-well plate, and the plate was placed back in an incubator (37°C, 5% $CO_2$) to allow for gelation to occur. This procedure was repeated for the "Epithelial," "Transitory," and "Mesenchymal" conditions. Each tube yielded a sufficient volume for 16 wells, allowing for various drug conditions to be tested across a singular cell condition. A second tube was prepared for each condition to check for consistency in cell phenotype across separately prepared tubes (technical replicates). 1 h after incubation, 120 $\mu$L of phenol red free growth media containing DMSO (Epithelial condition) or OHT (Mesenchymal and Transitory conditions) was added to the wells. Importantly, the concentration of the DMSO and OHT in media was prepared such that the final concentration would be 0.05% and 500 nM, respectively, taking into consideration the volume of media (120 $\mu$L) and the gel volume (60 $\mu$L).

Cells were allowed to acclimate to the 3D microenvironment for 24 h prior to the application of additional drug perturbations. Briefly, after 24 h in 3D culture, media containing various drug components were added (additional 120 $\mu$L, final media volume including gel = 300 $\mu$L) to yield a final concentration of 0.05% DMSO (negative control), 4 nM Paclitaxel (Taxol), or 500 nM Gefitinib. After three days, media was then carefully aspirated and growth media containing drugs +/- OHT were then replenished, which was repeated every three days for the total duration of the experiment.

**Confocal Microscopy.** Cell cluster movements and corresponding matrix displacements were imaged on day 7 after drug perturbations using a Nikon Eclipse Ti fluorescence microscope with spinning-disk confocal head (Crest Optics X-light V2), with light-guide coupled solid state illumination system (Lumencor Spectra-X3), sCMOS camera (Andor Neo), 20x Plan Apo objective (NA 0.75), GFP/FITC Filter Set (Chroma 49002), TRITC/DSRed Filter Set (Chroma 49004). For the duration of time-lapse imaging, cells were maintained in a humidified environmental chamber at 37 °C, 5% $CO_2$. For matrix displacement measurements, NIS Elements was used for automated image acquisition with z-steps of 0.6 $\mu$m from the bottom of the well to a height of 75 $\mu$m under consistent exposure times, camera gain/gamma control, and aperture. Images were acquired at 4 h intervals over a large number of wells ($n \approx 48$ for each experiment) for a total of 16 h. In these experiments, cell cytoplasm was imaged in GFP and beads in the RFP channel. At the end of time-lapse imaging, a reference state for the gel was obtained by lysing the cells within the hydrogels via sodium dodecyl sulfate.

Briefly, the 96-well plate lid was carefully removed, while keeping the plate within the microscope holder. Next, 100$\mu$L of media was removed from each well. Then, 100 $\mu$L of freshly prepared and warmed 4% sodium dodecyl sulfate in distilled, deionized water was added to each well to lyse cells. Complete lysis and relaxation of the gels can take several hours due to the initial diffusion into the gels, and variability in cluster sizes across conditions. As such, several additional z-stacks were acquired over a time-loop to ensure that the reference image used for displacement measurements used was a stabilized, relaxed gel. Due to the computational expense associated with analyzing the displacement fields for individual clusters and high N, we chose a representative timepoint (t=2) compared to the reference state of the relaxed gel > 12 h later.

**3D Cell-induced Deformation Measurement via T-PT.** Topology-based particle tracking reconstructed the cell-induced 3D displacement fields by tracking fluorescent polystyrene microspheres (1 $\mu$m) embedded as trackers particles in the silk-collagen matrix. The combination of the spinning disk confocal microscopy and the use of the low numerical aperture air-objective for high-throughput imaging of the 96-well plate setup resulted in low signal to noise volumetric images. The particle detection routine of the T-PT was modified with custom steps to account for the high noise in the volumes. First, the volume intensity along the $z$-direction was normalized because the particle fluorescence intensity rapidly dropped off when the confocal stacks deeper inside the ECM sample were imaged. The global image thresholding based particle detection could not account for such drastic variation in the local image intensity. Thus, the image intensity along the $z$-direction was normalized. For the image intensity normalization, the average image intensity along the $z$-axis was computed and then fit using a power series. Subsequently, the image intensity along the $z$-axis was normalized by multiplying raw image intensity with the inverse of the predicted average intensity from the power series as,

$$I_{norm}(x,y,z) = \frac{I(x,y,z)}{P(z)}. \tag{1}$$

The intensity normalized images ($I_{norm}$) were then spatially filtered with a 3D bandpass filter to suppress all the features of size lower or bigger than the characteristic size of the particles in the images. As a result, only the features retaining information of the characteristic size of the tracer particles remained in the images. The images were then filleted with a 3D Gaussian kernel with a sigma of 0.75. Subsequently, image thresholding was combined with watershed based segmentation method to identify connected components (simply connected white voxels in the binary image) in the images. The connected components were filtered according the their size corresponding to that of a single particle to uniquely identify a connected component corresponding to a single particle. Each filtered connected component had a minimum and maximum of 25 and 230 voxels respectively.

The coarse particle centers were estimated by computing the centroid of each connected components in the binarized images. Following, the rest of the particle localization and tracking routines of the T-PT remained unchanged with default tracking parameters, as described elsewhere (6).

**Cell Cluster Surface Segmentation.** The 3D cell cluster surface was segmented from the fluorescently labeled cytoplasm volumetric images (GFP channel). As a first step, the raw volumetric images were filtered using a median filter with $3 \times 3 \times 3$ voxel window to remove salt and pepper noise. Following the images were filtered with a 3D Gaussian filter with a sigma of 2.5. The images were then binarized using adaptive image thresholding based on the local mean intensity (first-order statistics) in the neighborhood of each voxel. The sensitivity for the adaptive thresholding was manually set for each image to segment the cell clusters from the background appropriately. From the binary images, the small connected components having a total number of voxels less than 8000 voxels were set to an intensity value of 0 in the binary images. Morphological operations were performed to remove holes in the binary images (7). The volumes of the segmented binary images were increased by 1.6 $\mu$m through a distance transform. Moreover, all the voxels in the lowermost and uppermost 4.8 $\mu$m (8 z-slices) of the imaging volume were set to 0 in the binarized images in order to correct for the increased noise levels in these locations. The 3D triangulated cell cluster surface was computed from the binary images using MATLAB's isosurface estimation at a target voxel value of 0.5. The triangulated cell cluster surface mesh was smoothed using accurate curvature flow smoothing (8).

**Data Registration.** The data registration was performed from the reconstructed motion field generated from tracked particles. First, the rigid translation in the volumes was registered by subtracting the median particle displacements. The median particle displacement provided a good estimate for the rigid image translation because the cell induced-displacements were localized in a much smaller region than the image volume size. Since most particles in the images did not experience cell-induced motion, and their displacement was caused by the rigid drift and rotation in the volume. Thus, the central statistics on the displacement field provided a reasonably accurate estimation of the rigid translation in the images.

After the correction for the rigid translation, the rotation experienced by the images along the $z$-axis was corrected by first computing the median rotation angle for each particle about the $x - y$ center of the image. From the median rotation angle, the rotation drift for each particle was corrected by adding an equal and opposite rotation drift to each particle displacement.

Lastly, the images also had a rigid displacement drift in the $x$ and $y$ direction as a function of $z$ position in the images possibly caused by backlash error in the stage motor. The stage motion drift was estimated by fitting a smoothing spline individually to the $u_x$ and $u_y$ with respect to the $z$ position of the particle. Then the stage motion drift was corrected by subtracting the predicted motion drift at each particle location based on its $z$ position.

**Mean Deformation Metrics.** In the absence of the material properties of silk-collagen ECM, 3D deformation field generated by the cell cluster were first characterized using the mean deformation metrics (MDM) (9). Based on the continuum mechanics framework, the mean displacement gradient $\langle \nabla u \rangle$ for a cell cluster in the image is given by

$$\langle \nabla u \rangle := \frac{1}{\mathrm{vol}(V_o)} \int_{V_o} \nabla u \, dV. \tag{2}$$

Here, $\nabla u$ is the gradient of the displacement field at a point, $dV$ is an infinitesimal volume element of a cell cluster in the reference image $V_o$ and $\mathrm{vol}(V_o)$ is the volume of the entire cell cluster. From the mean displacement gradient $\langle \nabla u \rangle$, the mean deformation gradient, $\langle F \rangle$, was computed as,

$$\langle F \rangle = I + \langle \nabla u \rangle. \tag{3}$$

The mean deformation gradient tensor, $\langle F \rangle$, provided various useful metrics in understanding the deformation of the cell cluster. However in the current form, determining $\langle F \rangle$ required computing $\nabla u$ at each point within the cell cluster (Equation 2). The particle tracking framework only provided deformation of the ECM and not of the points within the cell cluster. Therefore, using 2 and 3 the $\langle F \rangle$ could not be computed. However, utilizing the divergence theorem, $\langle \nabla u \rangle$ was computed from the cell cluster surface displacement as,

$$\langle \nabla u \rangle := \frac{1}{\mathrm{vol}(V_o)} \int_{\partial V_o} u \otimes n \, dA. \tag{4}$$

Here, $\partial V_o$ is the surface of the cell cluster, $dA$ is an infinitesimal element of $\partial V_o$ and $n$ is the unit vector normal to $dA$. Combining the results from Equation 4 and 3, the mean deformation gradient was computed. Once $\langle F \rangle$ was determined, various other useful metrics describing the cell cluster deformation like mean compressibility, contractility, and rotation of the cell cluster were computed. The mean compressiblity is the measure of volume change of the cell cluster and was determined as,

$$J = \det(\langle F \rangle). \quad [5]$$

The mean right stretch tensor, $\langle U \rangle$, was computed from the mean deformation gradient $\langle F \rangle$ through the relationship

$$\langle U \rangle = \left( \langle F \rangle^T \langle F \rangle \right)^{\frac{1}{2}}. \quad [6]$$

Subsequently, the mean rotation tensor was computed as,

$$\langle R \rangle = \langle F \rangle \langle U \rangle^{-1}. \quad [7]$$

From the mean right stretch tensor, the contractility of the cell cluster along principal directions was computed by determining the eigenvalues of $\langle U \rangle$. The mean rotation of the cell cluster was determined by computing the trace of the mean rotation tensor $\langle F \rangle$.

Essentially, once $\nabla u$ was determined, the computation of the remaining mean deformation metrics as straightforward. The $\langle \nabla u \rangle$ was numerically estimated from the relationship in the Equation 4 as

$$\langle \nabla u \rangle := \frac{1}{\mathrm{vol}(V_o)} \sum_{i=1}^{k} u_i \otimes n_i A_i. \quad [8]$$

Here, $\mathrm{vol}(V_o)$ was determined as the volume enclosed by the cell cluster surface triangulation mesh, $n_i$ is the unit outward normal vector to the $i^{th}$ face of the triangulation, $u_i$ is interpolated displacement field at the center of the $i^{th}$ face of the triangulation, and $A_i$ is the area of the $i^{th}$ face of the surface triangulation. The displacement field from the tracked particles at scattered locations was interpolated to the center of each face of the cell cluster surface triangulation using a linear scattered interpolation scheme.

**Shape Anisotropy Metric.** A shape anisotropy metric was used to quantify the anisotropy of the cell cluster shape. Shape anisotropy was defined as the ratio of the maximum Feret diameter and equivalent diameter from the binary maximum intensity projection of the cell cluster intensity on the $x - y$ plane. The binary maximum intensity projection of the cell was generated by summing the binary cell mask image along the $z$ axis and thresholding it to a value of 1. The maximum Feret diameter was measured as the maximum distance between any two boundary points on the antipodal vertices of the convex hull of the simply connected region of the cell cluster in the projected binary image. Equivalent diameter was defined as the diameter of the circle with the same area as the simply connected region of the cell cluster in the projected binary image.

**Hardware for Computation.** The deformation determination and kinematic analysis for all the cell clusters were conducted using the computational resources and services at the Center for Computation and Visualization, Brown University.

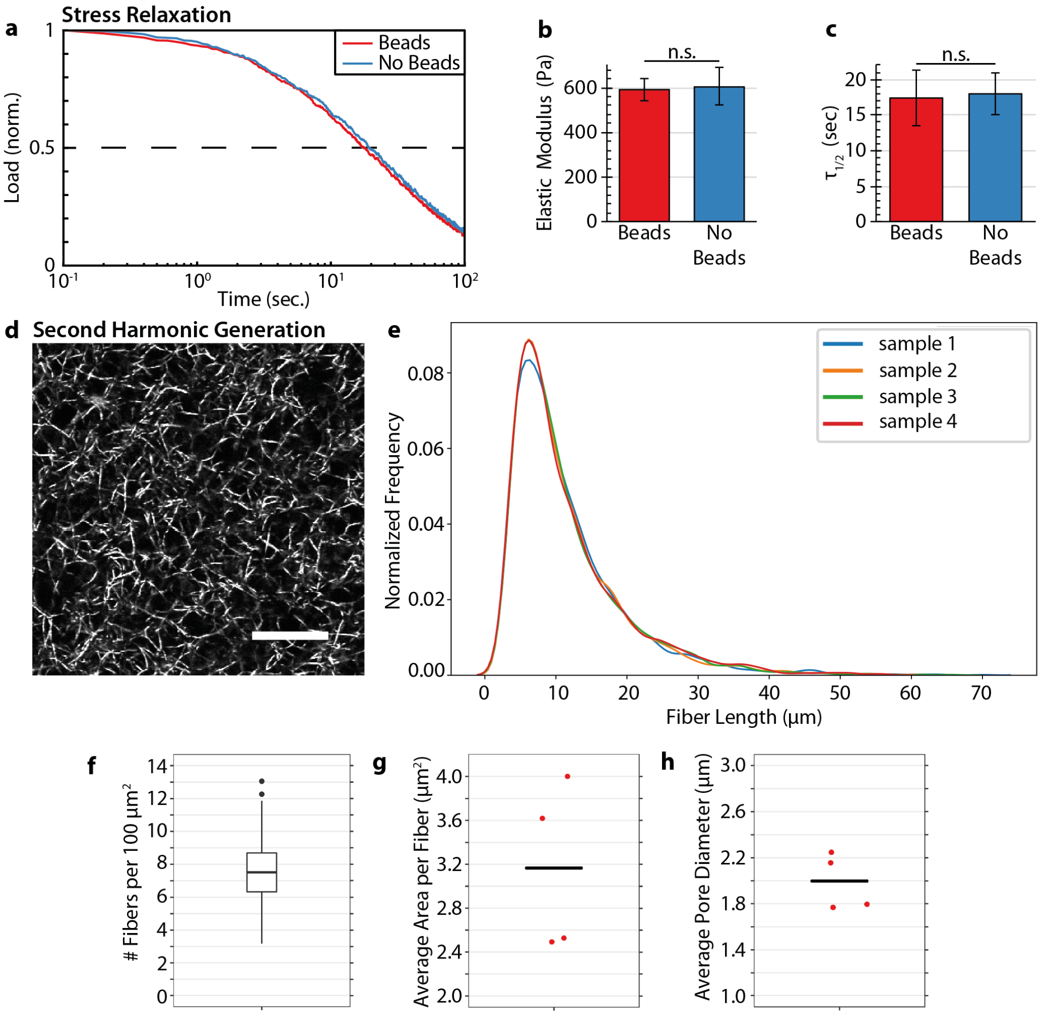


**Fig. S1.** **(a)** Representative stress-relaxation plot of 7.5 mg/mL silk fibroin and 1 mg/mL collagen I hydrogels prepared with and without tracer microparticles. Load is max-min normalized. **(b)** Comparison of elastic modulus for 7.5 mg/mL silk fibroin and 1 mg/mL collagen I hydrogels prepared with and without tracer microparticles. Barplot shows mean with error bars of one standard deviation. n.s. denotes no statistical significance by pair-wise Student's t-test. **(c)** Stress relaxation half-life $\tau_{1/2}$ for 7.5 mg/mL silk fibroin and 1 mg/mL collagen I hydrogels prepared with and without tracer microparticles. Barplot shows mean with error bars of one standard deviation. n.s. denotes no statistical significance by pair-wise Student's t-test. **(d)** Representative second harmonic generation imaging of fibrillar collagen I (1 mg/mL) in 7.5 mg/mL silk fibroin (not visible). **(e)** Statistical distribution of collagen fiber length in segmented fibers. **(f)** Statistical distribution of local fiber density, scaled per 100 $\mu m^2$ for ease of comparison. **(g)** Image-averaged pore area based on the area of a circle that fits within the square box that contains one fiber (averaged over the entire image). **(h)** Characteristic mesh size calculated as the square root of the pore area.

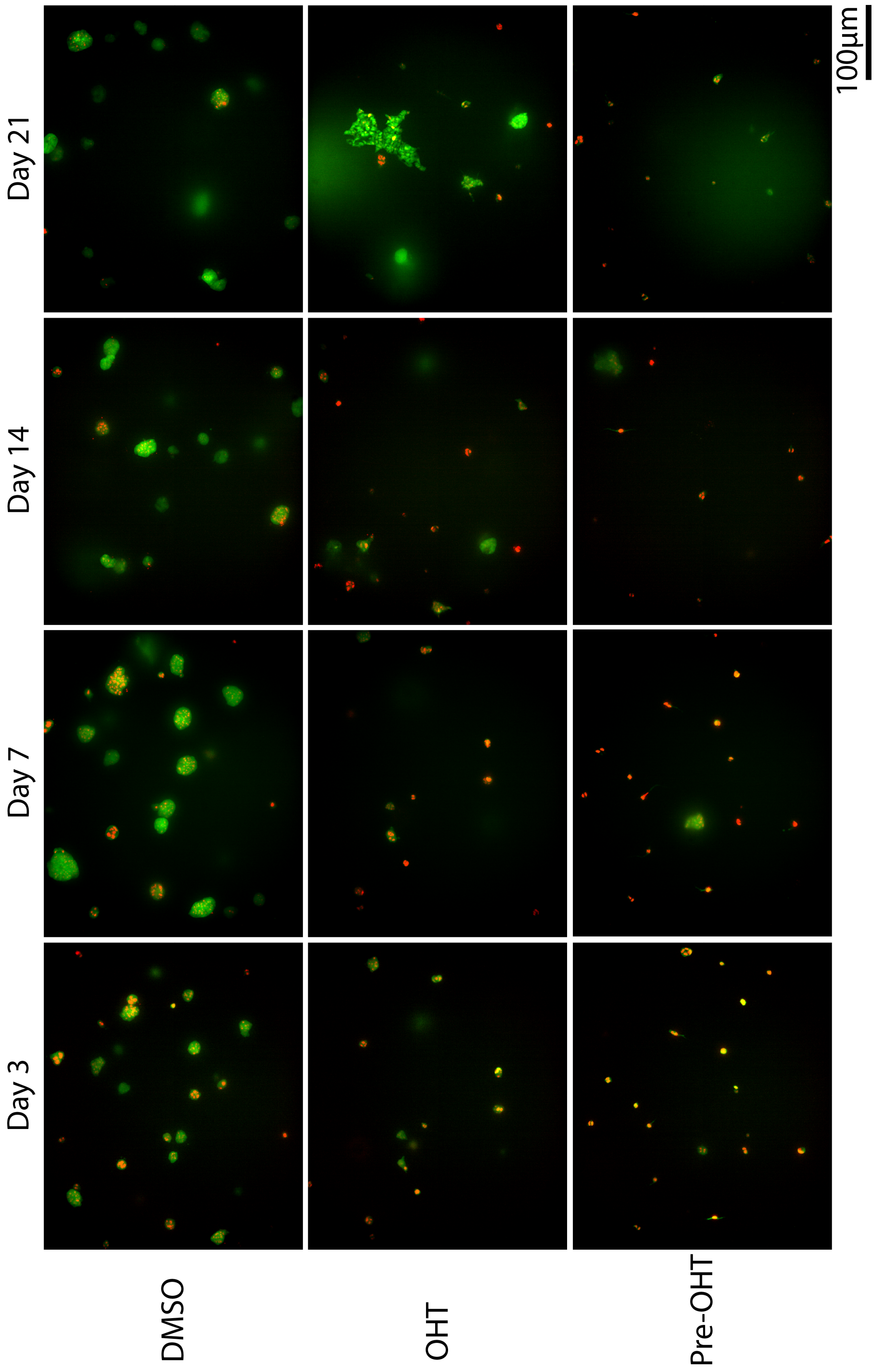


**Fig. S2.** Representative confocal z-slice images of MCF-10A cells cultured in 3D silk collagen hydrogels for varying durations (Days 3,7, 14, 21) and treatment conditions (DMSO and OHT treatments, OHT pre-treatment). Green (cytoplasmic GFP), red (nuclear H2B-mCherry)

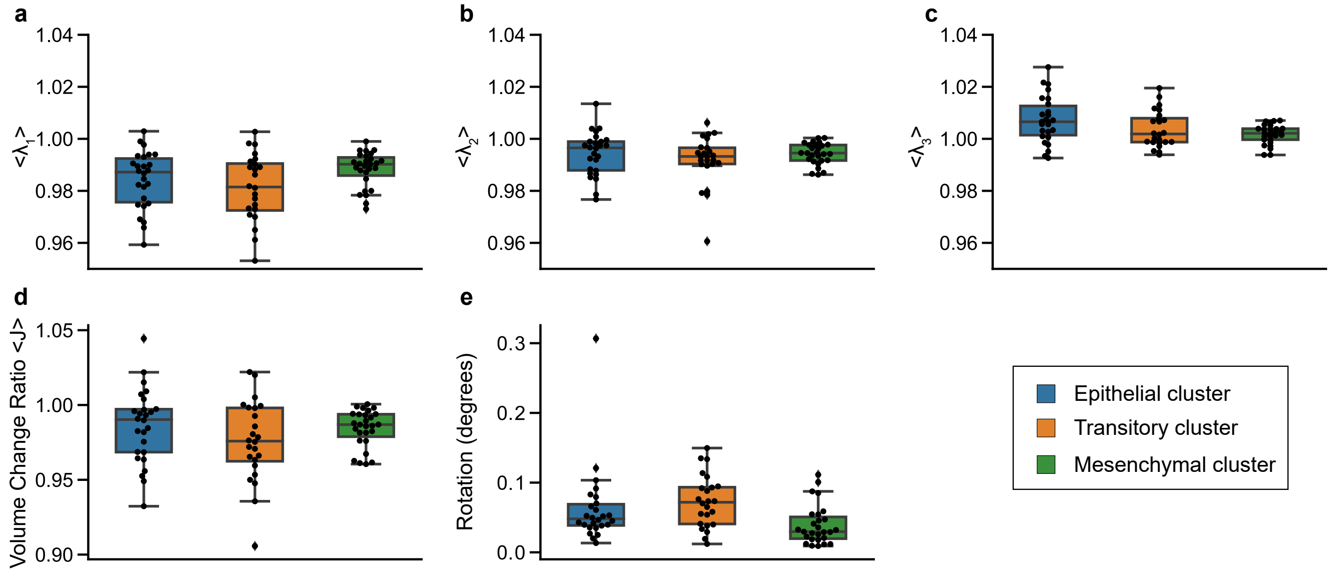


**Fig. S3.** Mean deformation metrics based on mean displacement gradient tensor $\langle \mathbf{u} \rangle$ for epithelial, transitory and mesenchymal phenotype. **(a,b,c)** Box plot of the three mean principal right stretch tensor eigenvalues values $\langle \lambda_i \rangle$ calculated from the mean right stretch tensor $\langle \mathbf{U} \rangle$. Positive value of $\langle \lambda_i \rangle$ represent mean expansion and the negative values of $\langle \lambda_i \rangle$ signify mean contraction of the cell along the corresponding principal axis. **(d)** Box plot of the mean volume change of the cell. A value of one indicates no change in volume. **(e)** Box plot of the mean rotation angle of the cell. The symbols ** represents $p$ values of $<0.01$.

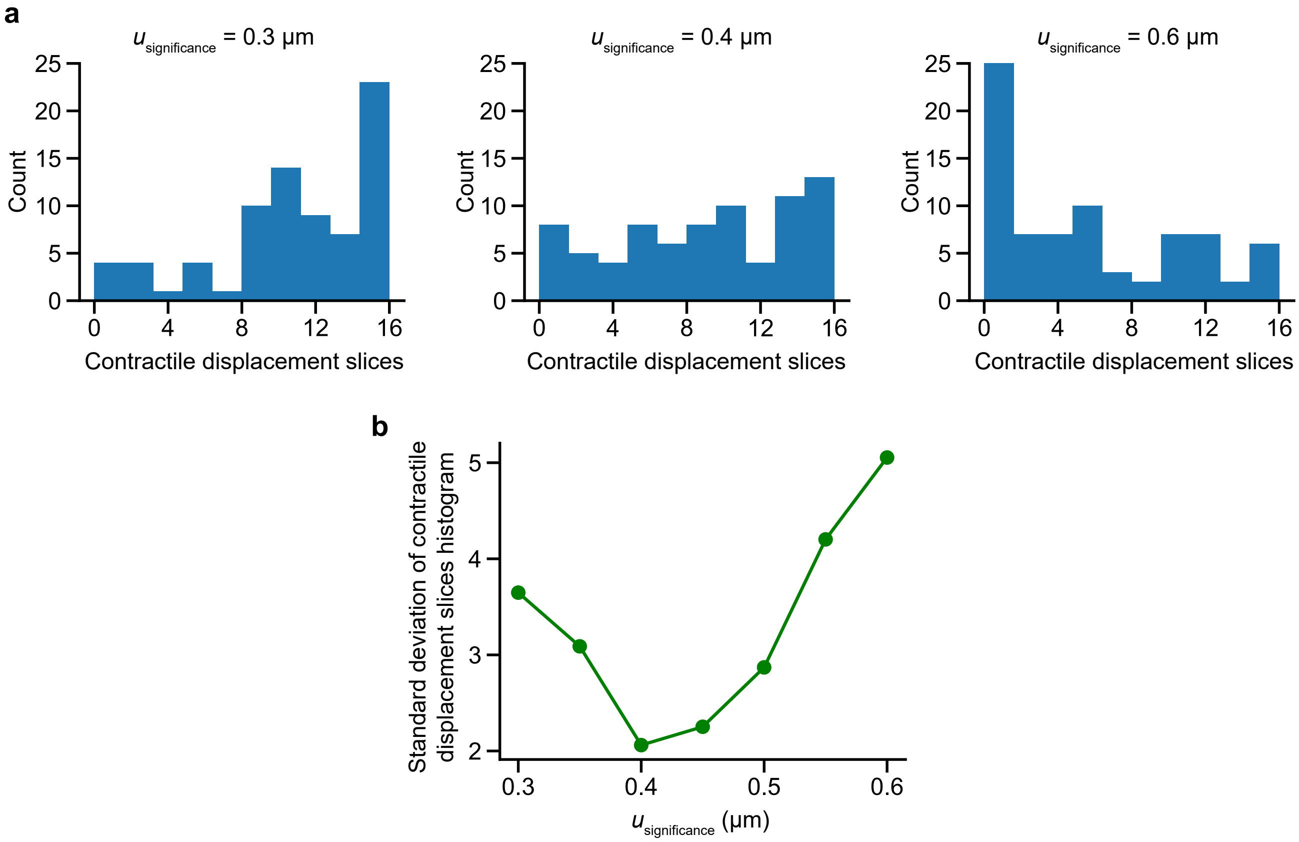


**Fig. S4.** Optimization to select an appropriate displacement threshold value, i.e., $u_{\text{significance}}$, for computing the DART metrics.The threshold value $u_{\text{significance}}$ was set to achieve a maximally even distribution in the number of contractile slices across all control clusters. This can be quantified by minimizing the standard deviation of contractile slices across the DART subvolumes, which achieves an optimal value at $u_{\text{significance}} = 0.4\mu m$. **(a)** Histograms of contractile displacement slices over a range of $u_{\text{significance}}$. **(b)** Plot of standard deviation of the histogram of contractile displacement slices versus $u_{\text{significance}}$ level.

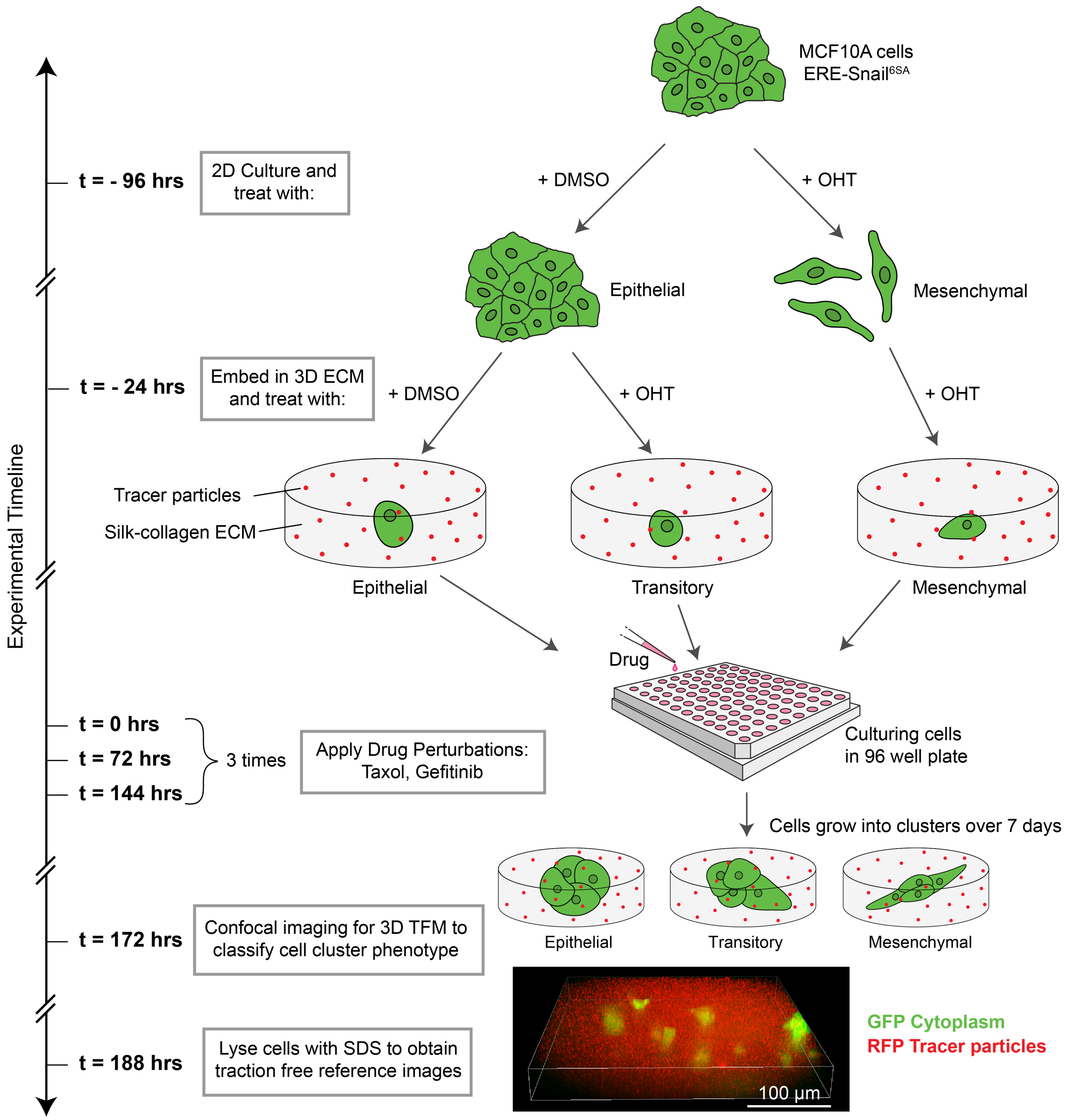


**Fig. S5.** Schematic illustrating experimental timeline.

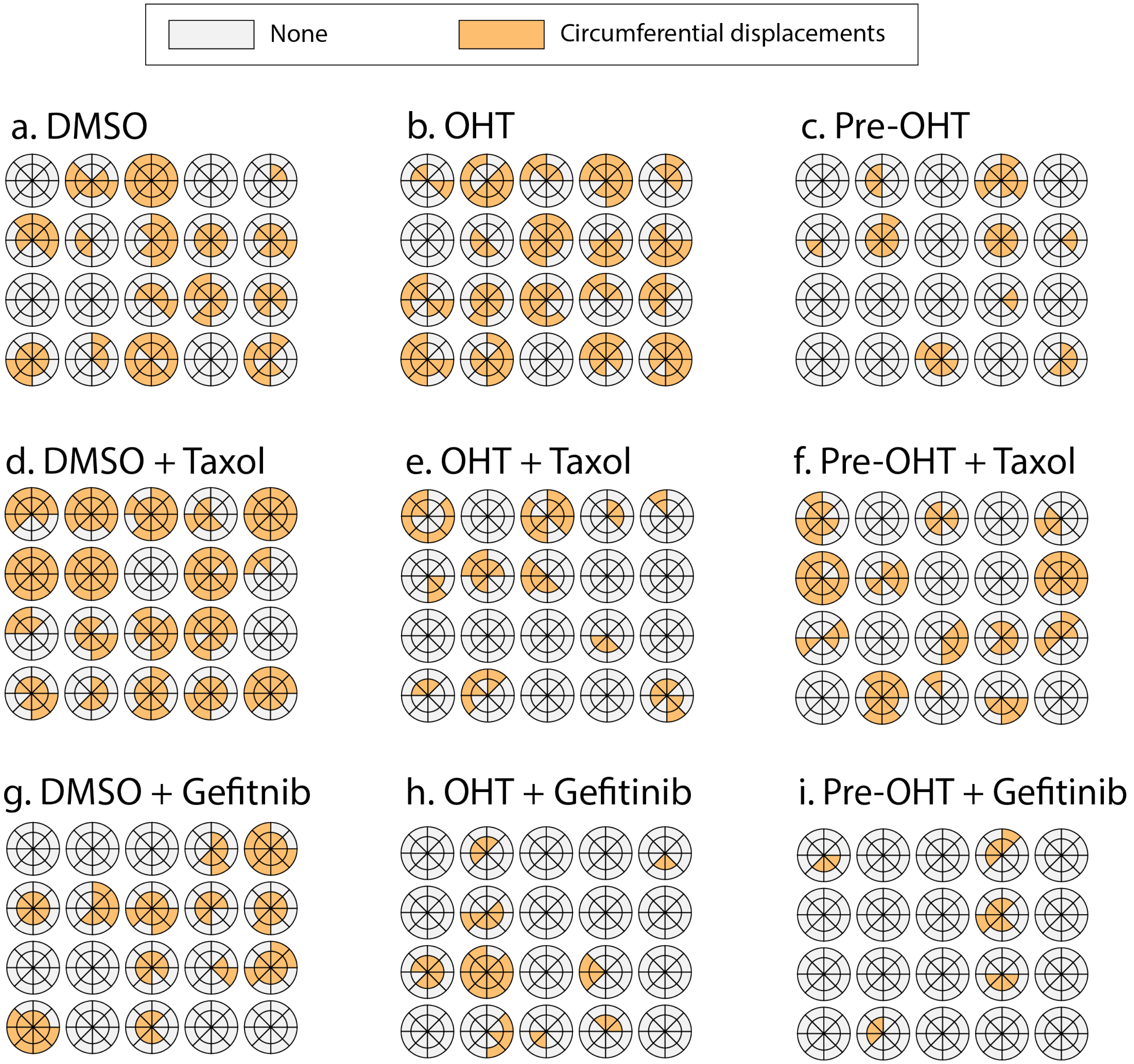


**Fig. S6.** Circumferential DARTS for twenty randomly selected clusters after DMSO treatment **(a)**, OHT treatment **(b)**, and OHT-pretreatment **(c)**. Circumferential DARTS for twenty randomly selected clusters with Taxol and DMSO treatment **(d)**, OHT treatment **(e)**, and OHT-pretreatment **(f)**. Circumferential DARTS for twenty randomly selected clusters with Gefitinib and DMSO treatment **(g)**, OHT treatment **(h)**, and OHT-pretreatment **(i)**.

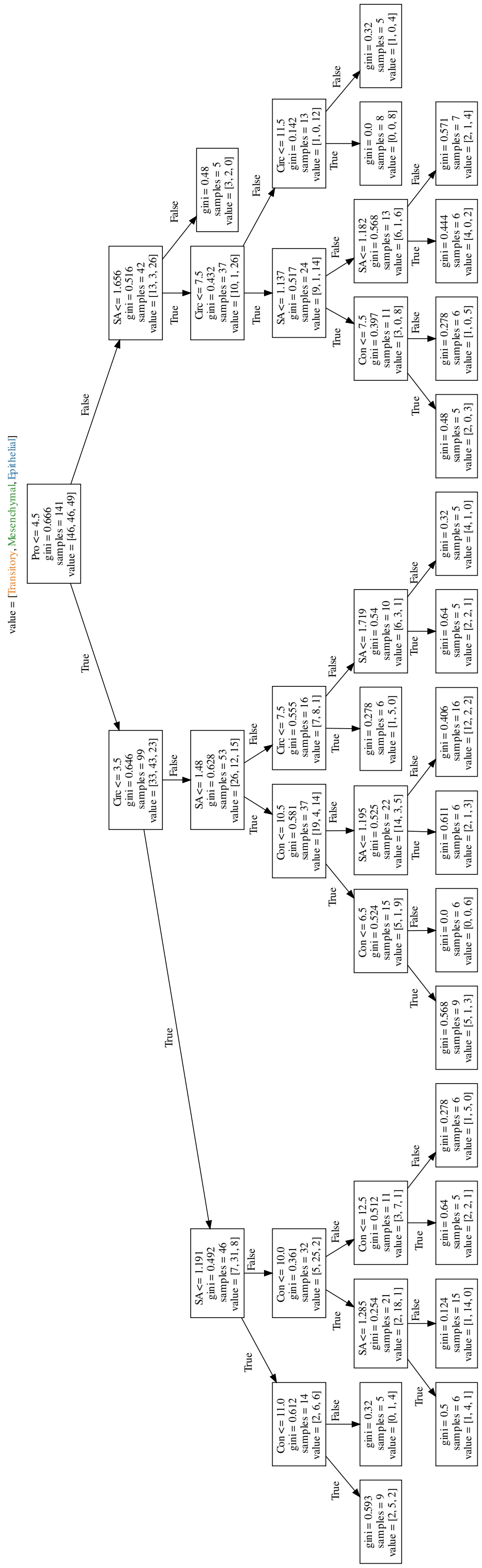


**Fig. S7.** Decision tree analysis for epithelial, transitory, and mesenchymal cluster mechanophenotypes. Briefly, clusters are classified as epithelial if protrusive slices > 4.5 unless shape anisotropy > 1.65 or circumferential slices < 7.5. Clusters are classified as transitory if protrusive slices < 4.5 and circumferential slices > 3.5, unless shape anisotropy > 1.48 or contractile slices < 10.5. Finally, clusters are classified as mesenchymal if protrusions < 4.5 and circumferential slices < 3.5, unless contractile slices > 11 or shape anistropy < 1.3.

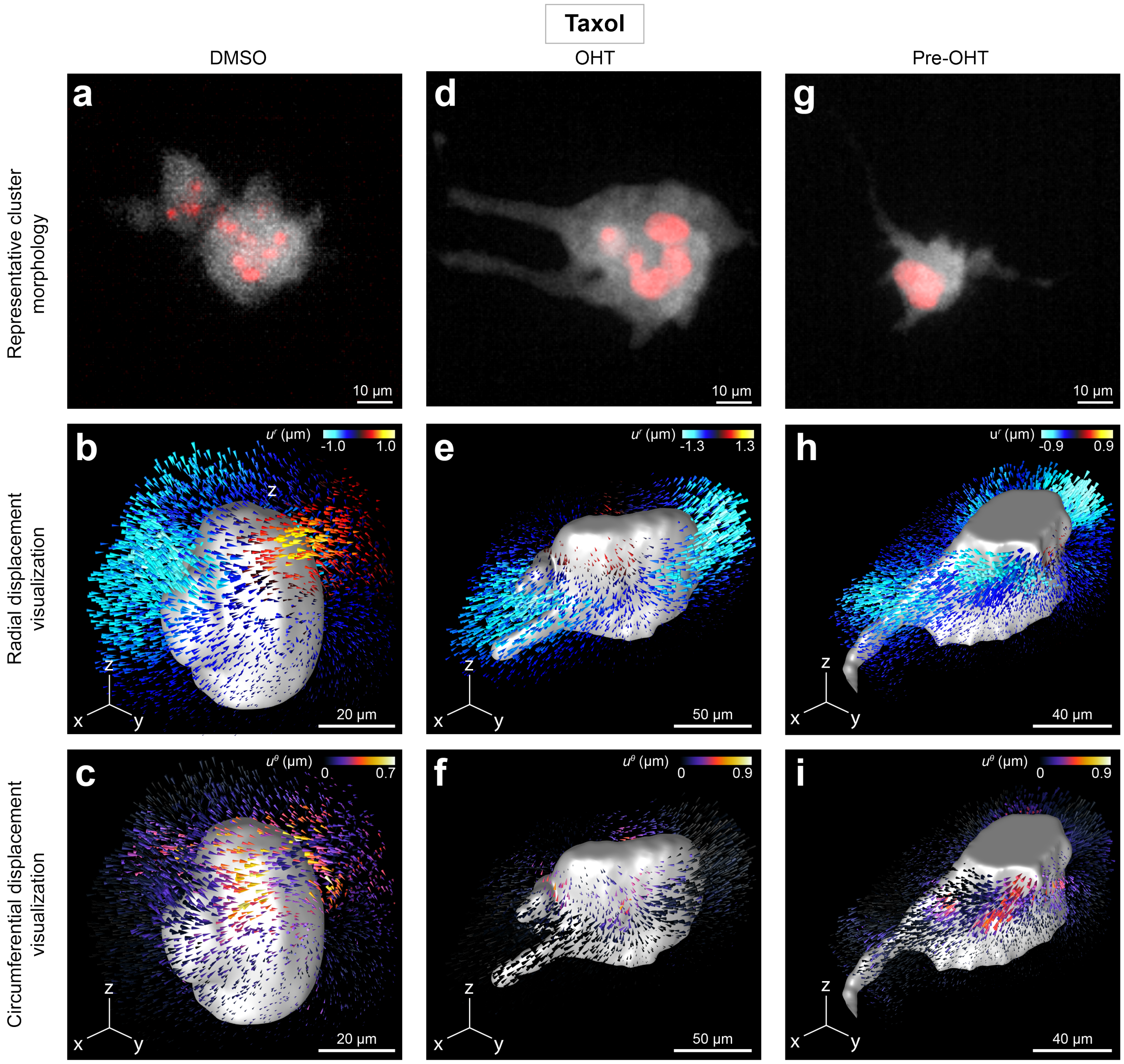


**Fig. S8.** Representative mechanophenotypes of clusters treated with 4 nM Taxol and DMSO **(a)**, OHT **(d)**, or pretreated with OHT **(g)**. Morphology of the cell clusters. The cytoplasm of the cell cluster is shown in gray (GFP) and the nucleus of the cells are shown in red (mCherry-H2B). **(b,e,h)** 3D radial displacements, $u^r$, about the center of a cell cluster shown for a representative epithelial, mesenchymal and transitory cell cluster. **(c,f,i)** Corresponding 3D circumferential displacements, $u^\theta$, about the center of a cell cluster shown for a representative epithelial, mesenchymal and transitory cell cluster.

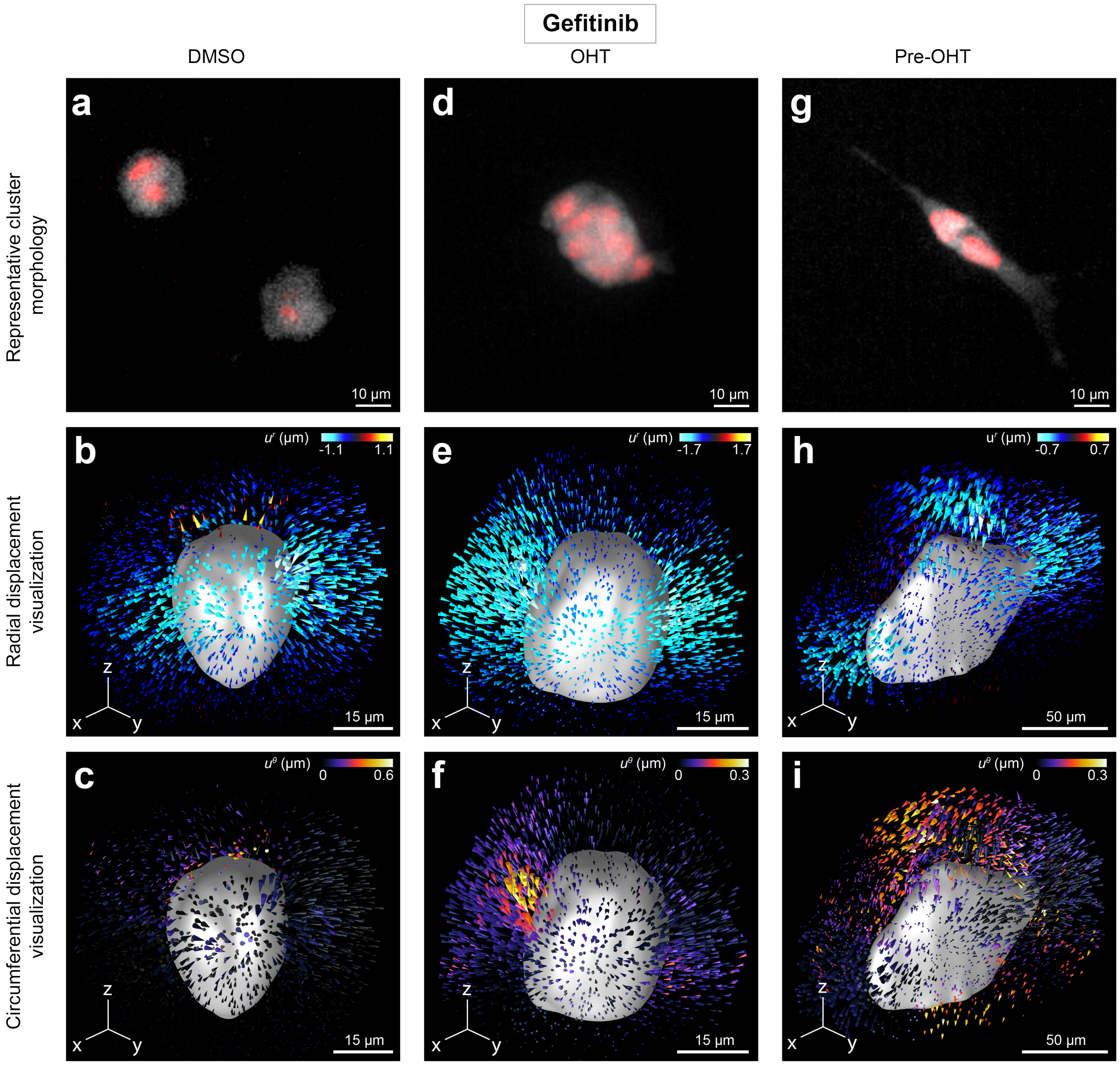


**Fig. S9.** Representative mechanophenotypes of clusters treated with 500 nM Gefitinib and DMSO **(a)**, OHT **(d)**, or pretreated with OHT **(g)**. **(a,b,c)** Morphology images for the cell clusters. The cytoplasm of the cell cluster is shown in green and the nucleus of the cells are shown in red. *Bottom left inset* shows the zoomed in cell cluster morphology for a representative cluster of within its class. *Top right* Schematic of representative shape and deformation associated with the phenotype with Gefitinib drug treatment. **(d,e,f)** 3D radial displacements, $u^r$, about the center of a cell cluster shown for a representative epithelial, mesenchymal and transitory cell clusters treated with Gefitinib. **(g,h,i)** 3D circumferential displacements, $u^\theta$, about the center of a cell cluster shown for a representative epithelial, mesenchymal and transitory cell clusters treated with Gefitinib.
